# Human brain organoids reveal accelerated development of cortical neuron classes as a shared feature of autism risk genes

**DOI:** 10.1101/2020.11.10.376509

**Authors:** Bruna Paulsen, Silvia Velasco, Amanda J. Kedaigle, Martina Pigoni, Giorgia Quadrato, Anthony Deo, Xian Adiconis, Ana Uzquiano, Kwanho Kim, Sean K. Simmons, Kalliopi Tsafou, Alex Albanese, Rafaela Sartore, Catherine Abbate, Ashley Tucewicz, Samantha Smith, Kwanghun Chung, Kasper Lage, Aviv Regev, Joshua Z. Levin, Paola Arlotta

## Abstract

Genetic risk for autism spectrum disorder (ASD) has been associated with hundreds of genes spanning a wide range of biological functions. The phenotypic alterations in the human brain resulting from mutations in ASD risk genes remain unclear, and the level at which these alterations converge on shared disease pathology is poorly understood. Here, we leveraged reproducible organoid models of the human cerebral cortex to identify cell type-specific developmental abnormalities associated with haploinsufficiency in three ASD risk genes, *SUV420H1* (*KMT5B*), *PTEN*, and *CHD8*. We performed comprehensive single-cell RNA-sequencing (scRNA-seq) of over 400,000 cells, and proteomic analysis on individual organoids sampled at different developmental stages to investigate phenotypic convergence among these genes. We find that within a defined period of early cortical development, each of the three mutations demonstrates accelerated development of cortical neurons. Notably, they do so by affecting different neuronal populations: excitatory deep layer (*SUV420H1*) and callosal (*PTEN*) neurons, and inhibitory interneurons (*CHD8*). This work shows that haploinsufficiency in ASD risk genes converge on early developmental defects in the generation of neurons of the cortical microcircuit.

## INTRODUCTION

Autism spectrum disorder (ASD) is a childhood-onset neurodevelopmental disorder characterized by cognitive, motor, and sensory deficits (Lord et al., 2020). ASD has a strong genetic component, with risk contribution from hundreds of genes (Grove et al., 2019, Rosenberg et al., 2009, Ruzzo et al., 2019, Sanders et al., 2012, Satterstrom et al., 2020). The shared developmental effects that cause this large and heterogeneous collection of genes to converge on the phenotypic features of ASD are poorly understood.

This is in part due to the lack of experimental models of human brain development, a process that largely occurs *in utero* and is therefore poorly accessible for study. The emergence of human brain organoids represents a major advance in modeling the cellular complexity of the developing human brain *in vitro* (Lancaster & Knoblich, 2014, Paşca, 2018). In combination with single-cell profiling technology, these models have the potential to enable the unbiased identification of the cell types, developmental events, and molecular mechanisms that are affected by risk-associated genes (Quadrato et al., 2016, Velasco et al., 2020).

Here we leveraged the use of reproducible organoid models of the cerebral cortex (Velasco et al., 2019a) to investigate the roles of three ASD risk genes in human cortical development. We chose *SUV420H1*, a histone methyltransferase (Jørgensen et al., 2013, Wickramasekara & Stessman, 2019) whose function in brain development is largely unknown but has been linked to neuroectodermal differentiation (Nicetto et al., 2013) and regulation of adult neural stem progenitor cell pools (Rhodes et al., 2016); *CHD8*, an ATP-dependent chromatin-remodeling factor associated with regulation of cell cycle and neuronal differentiation (Barnard et al., 2015, Breuss & Gleeson, 2016, Wade et al., 2019); and *PTEN*, which regulates cell growth and survival (Lee et al., 2018) and is linked with abnormal neuronal and glial development in animal models (Skelton et al., 2019). These three genes have emerged repeatedly as top hits in studies of ASD genetic risk (De Rubeis et al., 2014, Sanders et al., 2015, Satterstrom et al., 2020, Stessman et al., 2017, Yuen et al., 2017). They are expressed from very early stages of brain development, and homozygous-null disruption leads to early development lethality in animal models (Di Cristofano et al., 1998, Nishiyama et al., 2004, Schotta et al., 2008). All three are associated with severe neurodevelopmental abnormalities in humans, including shared phenotypes of high frequency of macrocephaly (Bernier et al., 2014, Faundes et al., 2018, Stessman et al., 2017, Yehia et al., 2020) suggestive of some degree of convergence in effect. However, the broad hypothetical functions and widespread expression of these genes make it difficult to predict how their mutation leads to disease and whether shared mechanisms may play a role.

By applying high-throughput single cell RNA-seq (scRNA-seq) and quantitative multiplexed proteomics of individual organoids carrying engineered haploinsufficient mutations in the *SUV420H1*, *CHD8*, or *PTEN* genes and their isogenic controls, we show that mutations in these genes converge on a shared effect of accelerated neurogenesis, which impacts distinct classes of cortical neurons in each mutation and acts through divergent molecular mechanisms. This work uncovers a neurobiological basis for how these genes may contribute to ASD pathology and identifies neurodevelopmental processes convergent across ASD risk genes.

## RESULTS

### Single-cell map of developing cortical organoids reveals cell type-specific enrichment of ASD risk genes

In order to determine whether human brain organoids could be used as models to understand the roles of ASD risk genes in human brain development, we first sought to verify expression of a large set of known risk genes in this cellular system across a developmental timeline. We generated a large dataset of scRNA-seq experiments from individual organoids produced from two different human induced pluripotent stem cell (iPSC) lines, GM08330 and Mito 210, using a recently-developed cortical organoid model (Figure 1, Supplementary Figure 1 and Supplementary Figure 2) (Velasco et al., 2019a).

**Figure 1.**
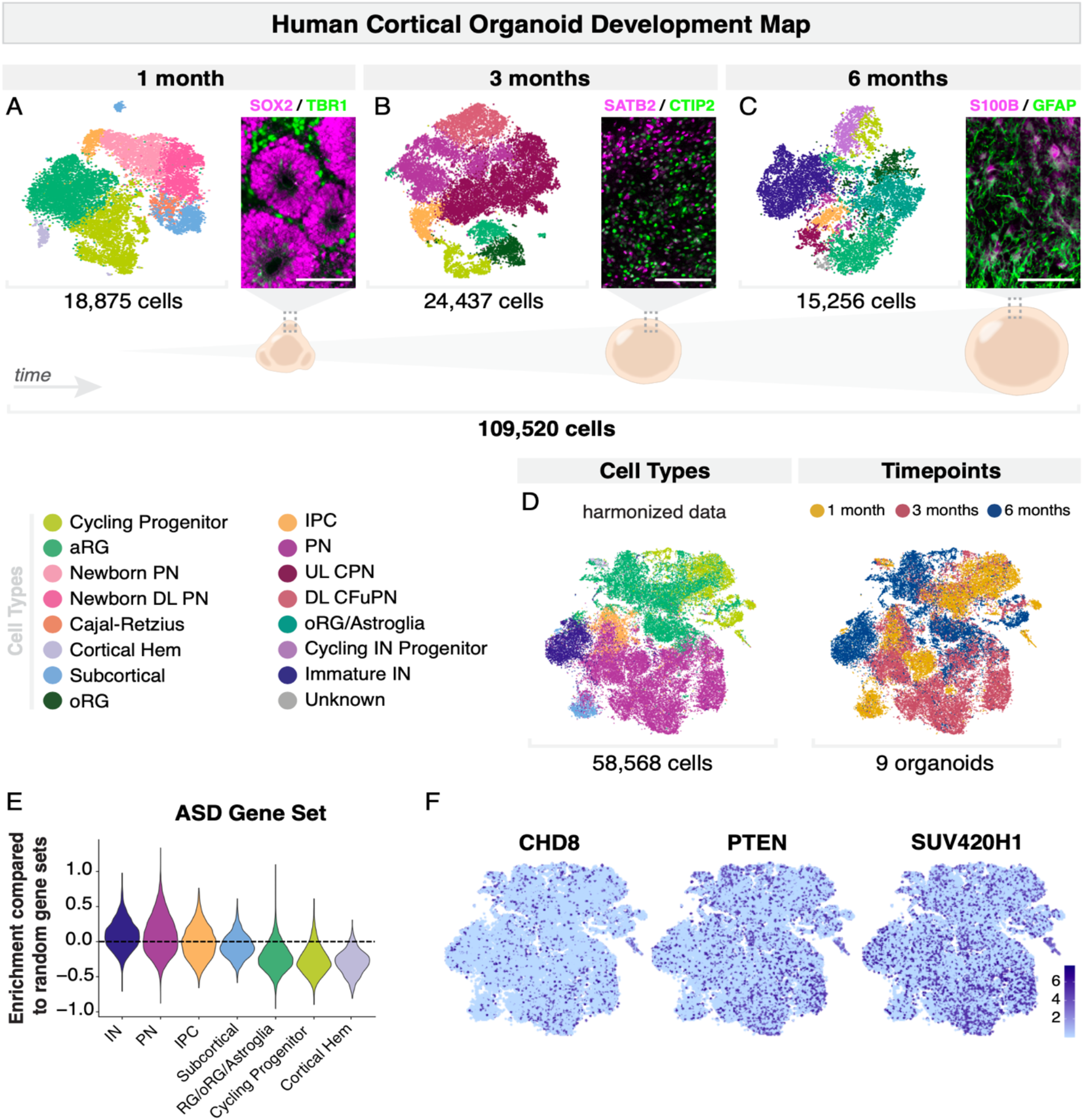
A Developmental map of human cortical organoids shows enrichment of ASD risk genes in neurons. A-C, scRNA-seq and immunohistochemistry analysis of organoids cultured for one, three and six months in vitro. Left, t-SNE plots (n = 3 organoids per timepoint, co-clustered). Cells are colored by cell type, and the number of cells per plot is indicated. Right, immunohistochemistry of specific markers. SOX2, a marker of neural progenitors and TBR1, a postmitotic neuronal marker, are shown at one month (A). SATB2, a marker of callosal projection neurons, and CTIP2, a marker of corticofugal projection neurons, are shown at three months (B). The astroglia markers S100B and GFAP are shown at six months (C). Scale bars are 100 μm. D, t-distributed stochastic neighbor embedding (t-SNE) plots of 58,568 cells from 9 organoids from the GM08330 cell line, at one, three and six months, after Harmony batch correction. Cells are colored according to cell type (left) and timepoint (right). E, Gene set expression scores for a set of 102 genes associated with ASD risk(Satterstrom et al., 2020) across cell types, in cells from d. Scores above 0 indicate enriched expression over similar sets of randomly chosen genes. F, t-SNE plots showing normalized expression of CHD8, PTEN, and SUV420H1 the three genes analyzed in this study, across the cells in d. PN, projection neurons; aRG, apical radial glia; UL CPN, upper layer callosal projection neurons; oRG, outer radial glia; DL CFuPN, deep layer corticofugal projection neurons; DL, deep layers; IPC, intermediate progenitor cells; IN, interneurons; t-SNE, t-distributed stochastic neighbor embedding.

We profiled individual organoids by immunohistochemistry and scRNA-seq across three stages of *in vitro* development: one month, when organoids contain a large proportion of progenitors and neurogenesis is beginning; three months, when progenitor diversity increases, including outer radial glia (oRG), and multiple subtypes of cortical projection neurons (PN) emerge; and six months, when interneurons (IN) and astroglia are present (Figure 1A-C and Supplementary Figure 1). At all timepoints, the cell type composition was highly reproducible between individual organoids, irrespective of batch and stem cell line (Supplementary Figure 1B) (Velasco et al., 2019a).

We next examined the cell type-specific expression of ASD risk genes in these organoids, using a set of 102 genes identified as contributing to ASD risk from the largest-to-date ASD exome sequencing study (Satterstrom et al., 2020). We first evaluated the expression of these genes as a set in each cell type of the combined scRNA-seq datasets (Figure 1D and Supplementary Figure 2A, D). Consistent with published data for the fetal human brain (Satterstrom et al., 2020), we found that as a set, ASD risk genes were highly enriched in excitatory projection neurons and interneurons, compared to the other cell types present in the organoids. The result was consistent across both stem cell lines analyzed (Figure 1E and Supplementary Figure 2B, E). We next examined expression of individual genes; we found that all 102 genes were expressed, in patterns that varied by cell type and developmental stage (Figure 1F and Supplementary Figure 2C, F). These results validate these organoids as a reliable model to investigate cortical cell types and their developmental trajectories, and suggest that they can be used to discover neurodevelopmental phenotypes associated with ASD risk genes.

To begin to understand the molecular effects of ASD risk genes and whether they show convergent effects on human brain development, we chose three genes for further study. We began with the above panel of ASD risk genes (Satterstrom et al., 2020) and focused on those that were: a) more strongly associated with ASD than intellectual disability/developmental delay and other neurodevelopmental abnormalities (Satterstrom et al., 2020, Stessman et al., 2017), b) predominantly found as *de novo* protein-truncating variants, which are more likely to be associated with increased ASD liability (Iossifov et al., 2012, Satterstrom et al., 2020), and c) expressed early in brain development (Satterstrom et al., 2020), and thus likely to cause developmental abnormalities that can be modeled in organoids.

Based on these criteria, we chose *SUV420H1*, *CHD8*, and *PTEN*, three genes that span different degrees of ASD risk (Satterstrom et al., 2020). In our scRNA-seq dataset, each of these three genes was expressed across multiple time points and cell types, precluding *a priori* prediction of cell type-specific effects (Figure 1F and Supplementary Figure 2C, F). We therefore selected them for comprehensive investigation using our organoid model system.

### *SUV420H1* haploinsufficiency causes increased production of deep layer cortical neurons

To investigate the effects induced by *SUV420H1* haploinsufficiency on human cortical development, we used CRISPR-Cas9 to generate a human iPSC line with a protein-truncating frameshift in the N-terminal domain of the SUV420H1 protein. After validation of the line, we produced cortical organoids from isogenic control and *SUV420H1* heterozygous mutant lines (Figure 2 and Supplementary Figure 3A-C). Immunohistochemistry for the progenitor markers EMX1 and SOX2, for the early pan-neuronal marker MAP2, and for the corticofugal projection neuron marker CTIP2, at one month in culture, showed that mutant organoids and their respective isogenic controls were able to efficiently begin neural development (Figure 2A, Supplementary Figure 3C). Notably, at 27 days *in vitro* (*d.i.v.*), *SUV420H1* mutant organoids were already substantially larger compared to controls, reminiscent of the macrocephaly phenotype observed in individuals carrying *de novo* mutations in the *SUV420H1* gene (n = 93 organoids, *p* = 0.0009, t-test; Figure 2B) (Faundes et al., 2018, Stessman et al., 2017).

**Figure 2.**
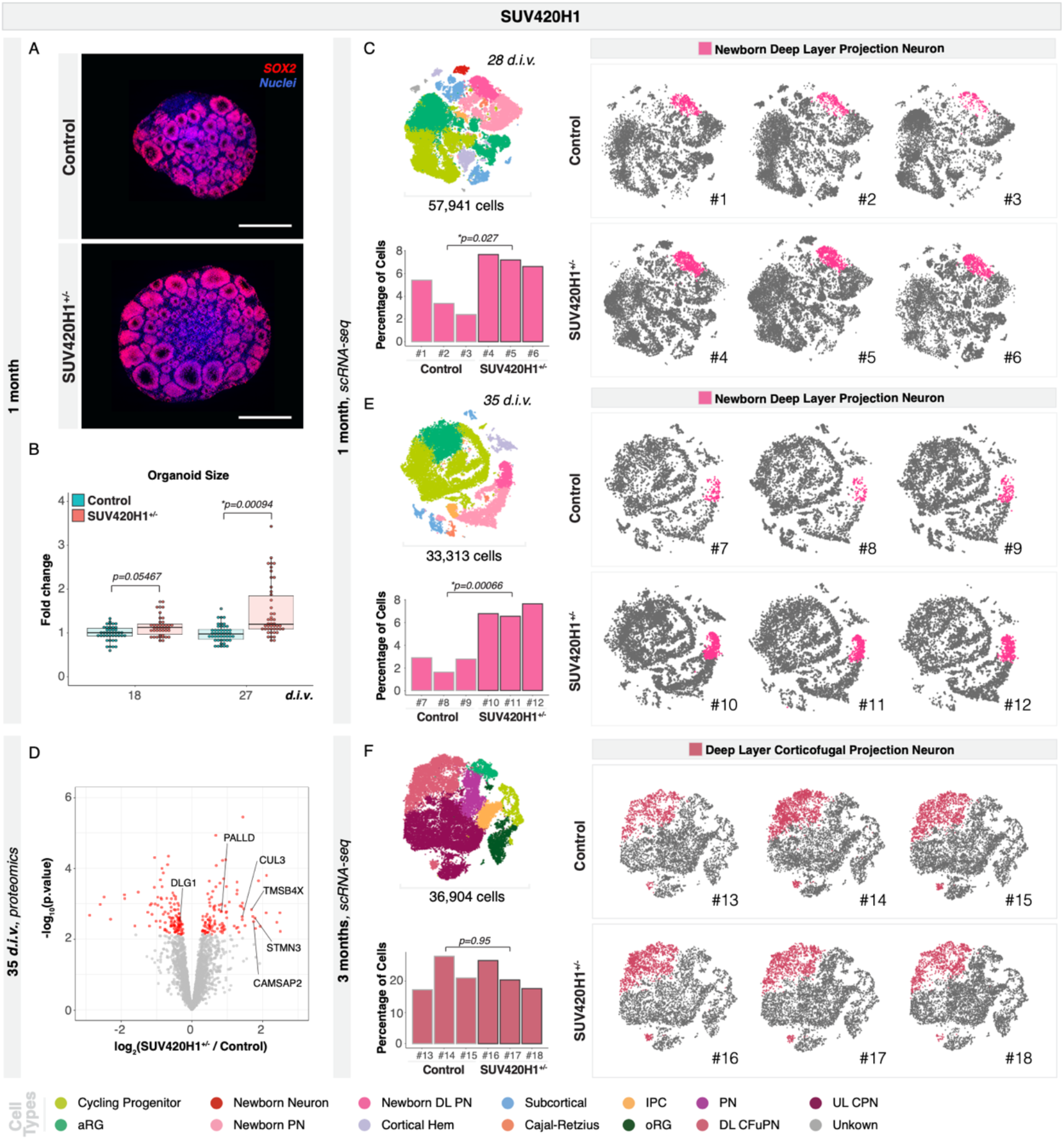
SUV420H1+/− mutant organoids show an increased number of deep layer projection neurons. A, Immunohistochemistry of SUV420H1^+/−^ mutant and control organoids cultured for 35 d.i.v. Optical section from the middle of whole-organoid dataset. Scale bars are 500 μm. SOX2, a marker of neuronal progenitors, is shown in red, and nuclei are shown in blue. B, Quantification of the size of control and SUV420H1^+/−^ organoids at 18 and 27 d.i.v. The ratio of organoid size compared to the average of control organoids in each batch is plotted. n = 42 for 18 d.i.v. control organoids, n = 42 for 18 d.i.v. mutant organoids, n = 47 for 27 d.i.v. control organoids, and n = 46 for 27 d.i.v. mutant organoids, from 4 experimental batches. P-values from a two-sided t test. C, E scRNA-seq data from 28 d.i.v. (C) and 35 d.i.v. (E) control and SUV420H1^+/−^ mutant organoids. Upper left shows combined t-SNE plots of control and mutant organoids (n = 3 single organoids per genotype). Cells are colored by cell type, and the number of cells per plot is indicated. Right, t-SNE plots for control and mutant individual organoids. Deep layer projection neuron populations are highlighted in color. Lower left, barcharts showing the percentage of cells for the highlighted cell populations in each control and mutant organoid. Adjusted p values for a difference in cell type proportions between control and mutant, based on logistic mixed models (see Methods) are shown. E, Volcano plot showing fold change versus adjusted p-value of measured proteins in MS experiments on SUV420H1^+/−^ versus control organoids cultured for 35 days (n = 4 single organoids per genotype). Significant DEPs are shown in red (adjusted p < 0.1). Proteins mentioned in the text are highlighted. F, scRNA-seq data from three month control and SUV420H1^+/−^ mutant organoids, as in C. d.i.v, days in vitro; PN, projection neurons; aRG, apical radial glia; UL CPN, upper layer callosal projection neurons; oRG, outer radial glia; DL CFuPN, deep layer corticofugal projection neurons; DL, deep layers; IPC, intermediate progenitor cells. DEPs: differentially expressed proteins; MS: mass spectrometry; t-SNE, t-distributed stochastic neighbor embedding.

We first focused our analysis on one month organoids. We performed transcriptional profiling by scRNA-seq of 57,941 single cells from individual control and mutant organoids at 28 *d.i.v.* (n = 3 single organoids per genotype; Figure 2C). The cellular composition of both control and mutant *SUV420H1* organoids at 28 *d.i.v.* was highly reproducible within genotypes and in each organoid. However, comparison between the genotypes showed a consistent increase in the cluster containing newborn deep layer projection neurons (the first-born neurons of the cortical plate (Greig et al., 2013, Lodato & Arlotta, 2015)), accompanied by a decrease in Cajal-Retzius cells (earlier-born pioneering neurons of the pre-plate (Marín-Padilla, 2015)) in the mutant organoids (Figure 2C and Supplementary Figure 3D). The newborn deep layer projection neurons, which expressed genes including *TBR1*, *FOXP2*, and *TLE4*, were consistently increased in every organoid analyzed (n = 3 single organoids per genotype; adjusted *p* = 0.027, logistic mixed models; Figure 2C).

To investigate the effect of *SUV420H1* haploinsufficiency on protein expression at one month in culture, we also performed whole-proteome analysis on single organoids from *SUV420H1* mutant and control lines, generated in a separate experimental batch (n = 3 single organoids per genotype), using isobaric tandem mass tag (TMT) multiplexed quantitative mass spectroscopy. We identified 233 proteins that were significantly differentially expressed (FDR < 0.1, moderated t-test; ≥ 4,000 proteins detected per sample, Figure 2D). Using Gene Set Enrichment Analysis (GSEA), we found that differentially expressed proteins (DEPs) were enriched for “neurogenesis”, as well as other broader processes such as “RNA binding” and “cell cycle” (Supplementary Figure 3E). DEPs included proteins associated with neuronal processes like axon guidance (e.g., PALLD, TMSB4X, CAMSAP2), synaptogenesis (e.g., DLG1, CUL3), and neuronal development (STMN3) (Figure 2D), suggesting an effect on neuronal processes.

We then investigated whether phenotypic abnormalities induced by *SUV420H1* heterozygosity persisted in more mature organoids. We performed scRNA-seq on two later timepoints, 35 *d.i.v*. and 3 months, using organoids derived from two new experimental batches (Figure 2E, F). In agreement with the increase in newborn deep layer projection neurons observed at 28 *d.i.v.,* scRNA-seq on 33,313 cells from *SUV420H1* control and mutant organoids sampled at 35 *d.i.v*, revealed an increase in this same population of cells, as well as smaller changes in intermediate progenitor cells, newborn projection neurons, and the subcortical clusters (Supplementary Figure 3D). The increase in deep layer projection neurons was more pronounced at this later developmental stage, and remained consistent in every individual organoid analyzed (n = 3 single organoids per genotype; adjusted *p* = 0.00066, logistic mixed models; Figure 2E). These data indicate that despite widespread *SUV420H1* expression (Figure 1F and Supplementary Figure 2C, F), its mutation results in a cell type-specific phenotype affecting a defined population of excitatory cortical neurons. Interestingly, this phenotype was rescued over developmental time in organoids; scRNA-seq on 36,904 cells from control and mutant organoids after three months *in vitro* showed that the percentage of deep layer corticofugal projection neurons was no longer significantly different between *SUV420H1* mutant and control (n = 3 single organoids per genotype; adjusted *p* = 0.95, logistic mixed models; Figure 2F), nor were any other cell types (Supplementary Figure 3D).

Altogether, these results indicate that deep layer projection neurons are generated earlier in *SUV420H1* mutant organoids compared to isogenic controls, and that this neuronal overproduction phenotype is restricted to a defined period during early development.

### *PTEN* haploinsufficiency leads to increased production of upper layer callosal projection neurons

To ascertain whether mutations in other ASD risk genes also result in altered production of early cortical neurons, we examined the molecular phenotype of *PTEN* haploinsufficiency in our organoid model. Using CRISPR-Cas9, we generated a new *PTEN* mutant iPSC line which we validated to carry a protein-truncating frameshift mutation in its phosphatase domain, a region found mutated in patients (Orrico et al., 2009), and produced mutant and isogenic control organoids (Figure 3A-E and Supplementary Figure 4A-C). Immunohistochemistry of one month organoids validated proper initiation of neural development in organoids (Supplementary Figure 4C). Resembling the *SUV420H1* mutants, *PTEN* haploinsufficient organoids also showed increased size compared to isogenic controls at 18 *d.i.v.* (n = 84 organoids, *p* = 2.39e-05, t-test; Figure 3A). This is in agreement with the macrocephaly phenotypes reported in patients with *PTEN* mutations (Yehia et al., 2020).

**Figure 3.**
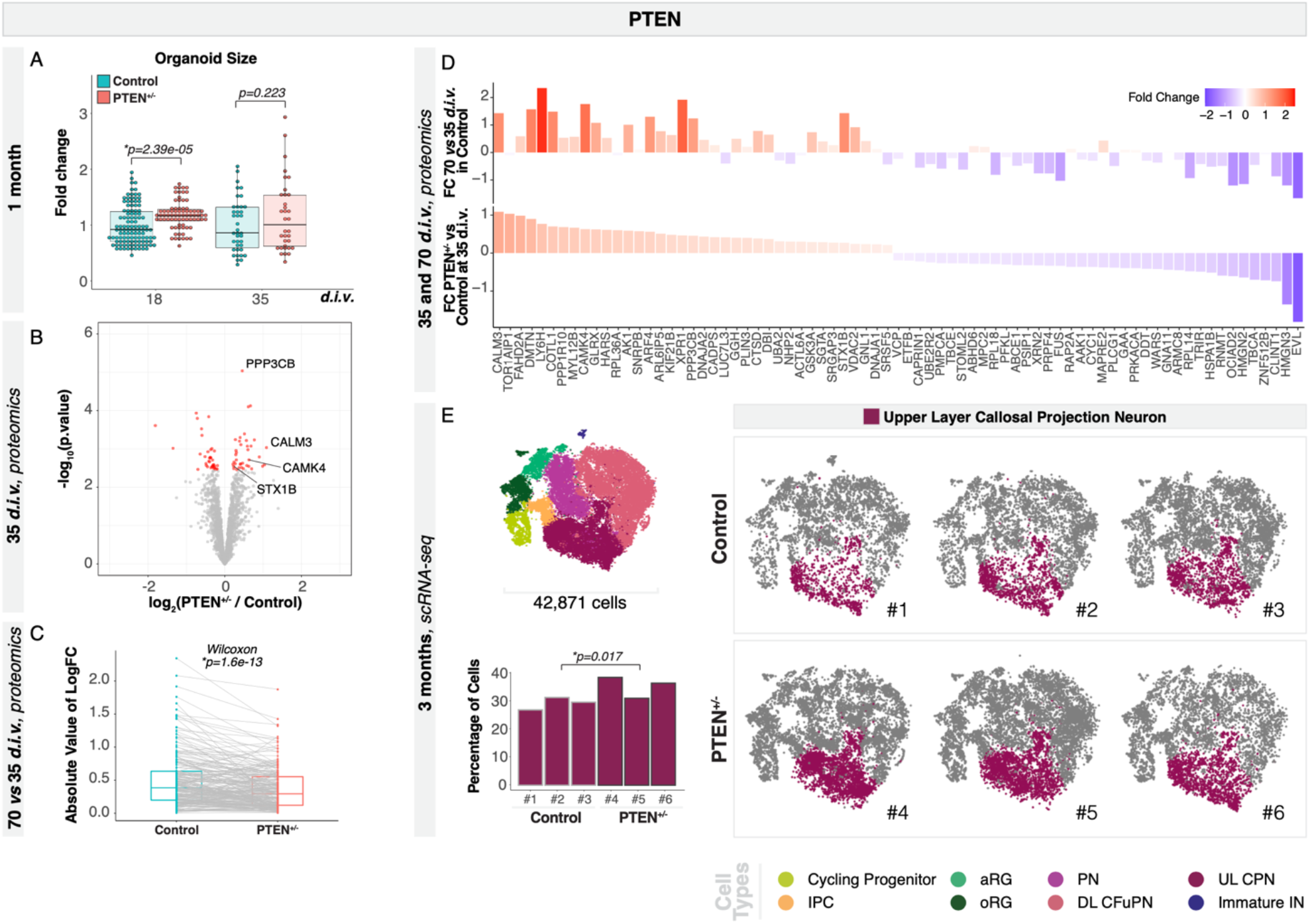
PTEN^+/−^ organoids show an increased number of callosal projection neurons. *A*, Quantification of the size of control and *PTEN*^+/−^ organoids at 18 and 35 *d.i.v*. The ratio of organoid size compared to the average of control organoids in each batch is plotted. n = 114 for 18 *d.i.v.* control organoids, n = 77 for 18 *d.i.v.* mutant organoids, n = 39 for 35 *d.i.v.* control organoids, and n = 34 for 35 *d.i.v.* mutant organoids, from 3 experimental batches. *P*-values from a two-sided t test. *B*, Volcano plot showing fold change versus adjusted *p*-value of measured proteins in MS experiments on PTEN^+/−^ versus control organoids cultured for 35 days (n = 4 single organoids per genotype). Significant DEPs are shown in red (adjusted *p* < 0.1). Proteins mentioned in the text are highlighted. *C*, Protein expression changes at 70 vs. 35 *d.i.v.* for control and mutant organoids. Gray lines connect values for the same protein in the two genotypes. *P*-value from a paired signed Wilcoxon rank test. Only significant DEPs are shown; for the same analysis done with all detected proteins see Supplementary Figure 4*H*. *D*, Comparison of protein expression changes in *PTEN*^+/−^ vs. control organoids at 35 *d.i.v.* (bottom) to changes in 70 vs. 35 *d.i.v.* control organoids (top). Color and y-axis indicate log_2_ fold change. Only proteins with significantly differential expression between control and mutant at 35 *d.i.v.* are shown. (n = 3 single organoids per genotype for 70 *d.i.v*). *E*, scRNA-seq data from *PTEN*^+/−^ and control organoids cultured for three months. Upper left shows combined t-SNE plots of control and mutant organoids (n = 3 single organoids per genotype). Cells are colored by cell type and the number of cells per plot is indicated. Right, t-SNE plots for control and mutant individual organoids. Callosal projection neuron populations are highlighted in color. Lower left, bar charts showing the percentage of callosal projection neurons in each control and mutant organoid. Adjusted p values for a difference in cell type proportions between control and mutant, based on logistic mixed models (see Methods) are shown. *d.i.v*, days *in vitro*; PN, projection neurons; aRG, apical radial glia; UL CPN, upper layer callosal projection neurons; oRG, outer radial glia; DL CFuPN, deep layer corticofugal projection neurons; DL, deep layers; IPC, intermediate progenitor cells; IN, interneurons. DEPs: differentially expressed proteins; MS: mass spectrometry; t-SNE, t-distributed stochastic neighbor embedding.

To investigate the cellular composition of *PTEN* mutant and isogenic control organoids, we performed scRNA-seq and proteomic analysis on individual organoids at one month in culture (Figure 3B and Supplementary Figure 4D). Analysis of 26,795 single cells at 35 *d.i.v.* showed that, in contrast to *SUV420H1*, in the *PTEN* mutant organoids there was no difference in the proportion of newborn deep layer projection neurons (n = 3 single organoids per genotype, adjusted *p* = 0.4, logistic mixed models; Supplementary Figure 4D, F) or any other cell type (Supplementary Figure 4F). In order to validate this result, we repeated the scRNA-seq experiment and included proteomic analysis using two new experimental batches. scRNA-seq of an additional 36,786 cells from a second batch of organoids grown for 35 *d.i.v* confirmed our first result, revealing no significant difference between genotypes in this replicate experiment (n = 3 single organoids per genotype, adjusted *p* = 0.56, logistic mixed models; Supplementary Figure 4E, F). However, whole-proteome mass spectrometry on single *PTEN* mutant and isogenic control organoids at one month in culture (n = 4 single organoids per genotype), showed significant differential expression of 75 proteins between genotypes (*p* < 0.1, moderated t-test; Figure 3B). GSEA identified “synapse” as the most enriched cellular localization, along with biological processes such as exocytosis (e.g., STX1B) and the calcium signaling pathway (e.g., PPP3CB, CALM3, and CAMK4), pointing to a potential effect of the *PTEN* mutation on neuronal development (Figure 3B and Supplementary Figure 4G).

We extended the whole-proteome analysis of *PTEN* mutant and isogenic control organoids to a later timepoint, 70 *d.i.v.* (n = 3 single organoids per genotype). While there were no differences in protein abundance between mutant and controls, when we examined the proteins that were differentially expressed between time points (70 *vs.* 35 *d.i.v.*) in each genotype, the mutant organoids showed significantly smaller expression changes over time than controls (*p* = 1.6e-13, Wilcoxin rank test, Figure 3C and Supplementary Figure 4H), indicating that the mutant 35 *d.i.v.* organoids resembled their older counterparts more closely than the control 35 *d.i.v.* organoids. Furthermore, the mutation-driven changes (i.e., the set of proteins that were differentially expressed between genotypes in the 35 *d.i.v.* organoids) were correlated with time-driven changes (i.e., the set of proteins that changed in both the control and mutant proteomes through time, 70 vs 35 *d.i.v.*), indicating that *PTEN* mutation affects the expression of developmentally-regulated proteins (R = 0.76, *p* < 1e-15, Pearson correlation; Figure 3D and Supplementary Figure 4I). This result points to a precocious development of *PTEN* mutant organoids.

We then performed scRNA-seq profiling of 42,871 additional cells from a later stage of *PTEN* mutant and control organoids, three months in culture (n = 3 separate organoids per genotype). Mutant organoids at this stage showed a small but significant increase in cycling progenitors and oRG (Supplementary Figure 4F), consistent with a prior study reporting that *PTEN* homozygous loss-of-function human cerebral organoids showed overgrowth coupled with an expansion of neural progenitors (Li et al., 2017). However, at this stage, the *PTEN* mutant organoids also showed a notable overproduction of a single population of excitatory neurons: callosal projection neurons (adjusted *p* = 0.017, logistic mixed models; Figure 3E and Supplementary Figure 4F). These results indicate that, while affecting a different population of excitatory neurons than the *SUV420H1* mutation, both *SUV420H1* and *PTEN* haploinsufficiency converge on a shared phenotype of imbalanced production of cortical excitatory neurons.

### *CHD8* haploinsufficiency causes overproduction of cortical interneurons

In order to determine if alterations in the tempo of neurogenesis of cortical neurons was shared by a mutation in a third gene, we obtained a human embryonic stem cell (ESC) line heterozygote for a protein-truncating frameshift in the helicase C-terminal domain of the *CHD8* gene, and generated mutant and isogenic control organoids (Figure 4A-D and Supplementary Figure 5A-C). Immunohistochemistry at one month *in vitro* indicated that haploinsufficiency in *CHD8* does not disturb proper initiation of neurodevelopment using this organoid protocol (Supplementary Figure 5C). Notably, as with *SUV420H1* and *PTEN*, *CHD8* mutant organoids showed increased size at 25 *d.i.v.* (n = 33 organoids, *p* = 2.305e-05, t-test; Figure 4A), consistent with macrocephalic phenotypes reported in 80% of patients with mutations in *CHD8* (Bernier et al., 2014).

**Figure 4.**
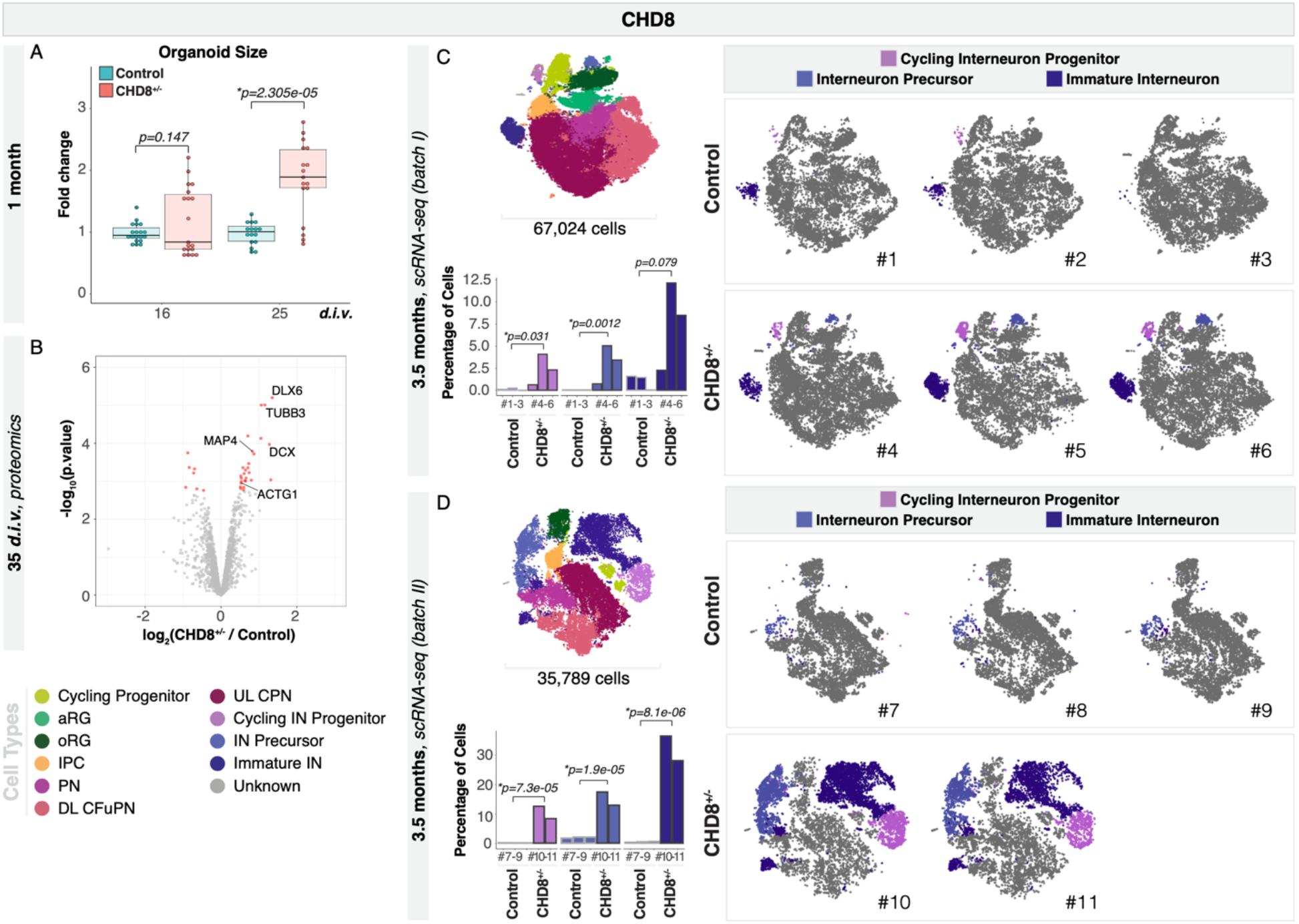
CHD8^+/−^ organoids show an increased number of interneurons. ***A***, Quantification of the size of control and *CHD8*^+/−^ organoids at 16 and 25 *d.i.v*. The ratio of organoid size compared to the average of control organoids in each batch is plotted. n = 19 for 16 *d.i.v.* control organoids, n = 19 for 16 *d.i.v.* mutant organoids, n = 16 for 25 *d.i.v.* control organoids, and n = 17 for 25 *d.i.v.* mutant organoids, from 2 experimental batches. *P*-values from two-sided t test. *B*, Volcano plot showing fold change versus adjusted *p*-value of measured proteins in MS experiments on *CHD8*^+/−^ vs. control organoids cultured for 35 days (n = 3 single organoids per genotype). Significantly DEPs are shown in red (adjusted *p* < 0.1). *C* & *D*, scRNA-seq data from *CHD8*^+/−^ and control organoids cultures for 3.5 months, from experimental batch I (*C*) and II (*D*). Upper left shows combined t-SNE plots of control and mutant organoids (n = 3 single organoids per genotype, except batch II mutant for which n = 2). Cells are colored by cell type and the number of cells per plot is indicated. Right, t-SNE plots for control and mutant individual organoids. Interneurons and their progenitor populations are highlighted in color. Lower left, bar charts showing the percentage of interneurons and their progenitors in each control and mutant organoid. Adjusted p values for a difference in cell type proportions between control and mutant, based on logistic mixed models (see Methods) are shown. *d.i.v*, days *in vitro*; PN, projection neurons; aRG, apical radial glia; UL CPN, upper layer callosal projection neurons; oRG, outer radial glia; DL CFuPN, deep layer corticofugal projection neurons; DL, deep layers; IPC, intermediate progenitor cells; IN, interneurons. DEPs: differentially expressed proteins; MS: mass spectrometry; t-SNE, t-distributed stochastic neighbor embedding.

We first performed whole-proteome mass spectrometry on one month *CHD*8 mutant and control organoids (n = 3 for each of mutant and control), and found significant differential expression of 35 proteins between genotypes. The interneuron fate regulator DLX6 (Lim et al., 2018) was found among the most upregulated proteins in the mutant (Figure 4B), suggesting an earlier commitment of mutant progenitors to an interneuron fate, as proposed by a prior study using bulk RNA-seq analysis of *CHD8* mutant human cerebral organoids (Wang et al., 2017). Additionally, proteins associated with neuronal differentiation and cytoskeleton remodeling (TUBB3, DCX, MAP4, ACTG1) were upregulated in mutant organoids (Figure 4B). Enriched biological processes in GSEA included “neuron differentiation”, “neuron projection”, and “neuron development” (Supplementary Figure 5D), suggesting that cortical neurons might be affected in *CHD8* haploinsufficient organoids.

These results prompted us to perform scRNA-seq analysis on *CHD8* mutant and control organoids cultured for 3.5 months, a stage when interneurons begin to appear (67,024 cells, n = 3 single organoids per genotype, 109 *d.i.v.*). At this stage, our analysis revealed a dramatically increased number of cycling interneuron progenitors, interneuron precursors, and immature interneurons in mutant organoids (cycling interneuron progenitors: adjusted *p* = 0.031, interneuron precursors: adjusted *p* = 0.0012, immature interneurons: adjusted *p* = 0.079, logistic mixed models; Figure 4C). We also observed a slight increase in the number of apical radial glia (Supplementary Figure 5E), suggesting that the increased number of progenitors might not be restricted to the ones committed to an interneuron fate.

We repeated this experiment organoids from a separate experimental batch, profiling 35,789 additional single cells from control and mutant organoids collected at 3.5 months (107 *d.i.v.)*. In this new experiment, we again observed a dramatic increase in the numbers of immature interneurons, as well as in the numbers of interneuron precursors and cycling interneuron progenitors (n = 3 single organoids per genotype; cycling interneuron progenitors: adjusted *p* = 7.3e-05, interneuron precursors: adjusted *p* = 1.9e-05, immature interneurons: adjusted *p* = 8.1e-06, logistic mixed models; Figure 4D). The increase in interneurons was so pronounced that the represented proportion of all the other cell types were reduced in the mutant organoids (Supplementary Figure 5E). These data demonstrate that, as observed for *SUV420H1* and *PTEN* mutations, *CHD8* haploinsufficiency is associated with precocious differentiation of a specific neuronal lineage of the cortical microcircuit, in this case, cortical interneurons.

Altogether, these data indicate that despite the widespread expression and broad functions of each of the three ASD risk genes analyzed here, they converge on overproduction of cortical neuronal lineages. Notably, each mutation affects different subsets of neuronal populations, indicating that ASD pathology in these three genes may converge on a broad phenotype of precocious neuronal overproduction, rather than on a shared cell type or molecular pathway.

### Developmental trajectories of *SUV420H1*, *PTEN*, and *CHD8* mutant organoids show accelerated neurogenesis of specific neuronal subtypes

The increased number of cortical neurons in the three mutants raised the question of whether overproduction of neurons was the result of accelerated differentiation. We therefore calculated pseudotime trajectories for each scRNA-seq dataset using Monocle3 (Cao et al., 2019). For all three mutants and their isogenic controls, the pseudotemporal ordering of cell types approximated that of *in vivo* human development, following a trajectory that progressed from progenitors to neuronal cell types (Figure 5A-F, Supplementary Figure 6A and Supplementary Figure 7A, B). For *SUV420H1* at 35 *d.i.v.*, the cells split into two major partitions, one containing progenitors traversing the cell cycle, and one containing cycling progenitors maturing towards newborn deep layer neurons (Supplementary Figure 6A). The latter contained the neuronal populations most consistently affected by the mutation, and was thus selected as the partition of interest. For *PTEN*, the cells were all assigned to one trajectory, with progenitors at the beginning, and a branch point between cells progressing to callosal projection neurons or corticofugal projection neurons (Supplementary Figure 7A). Finally, for *CHD8*, the partition of interest showed cells progressing from radial glia to interneurons, and a separate branch contained the progression from radial glia to projection neurons (Supplementary Figure 7B).

**Figure 5.**
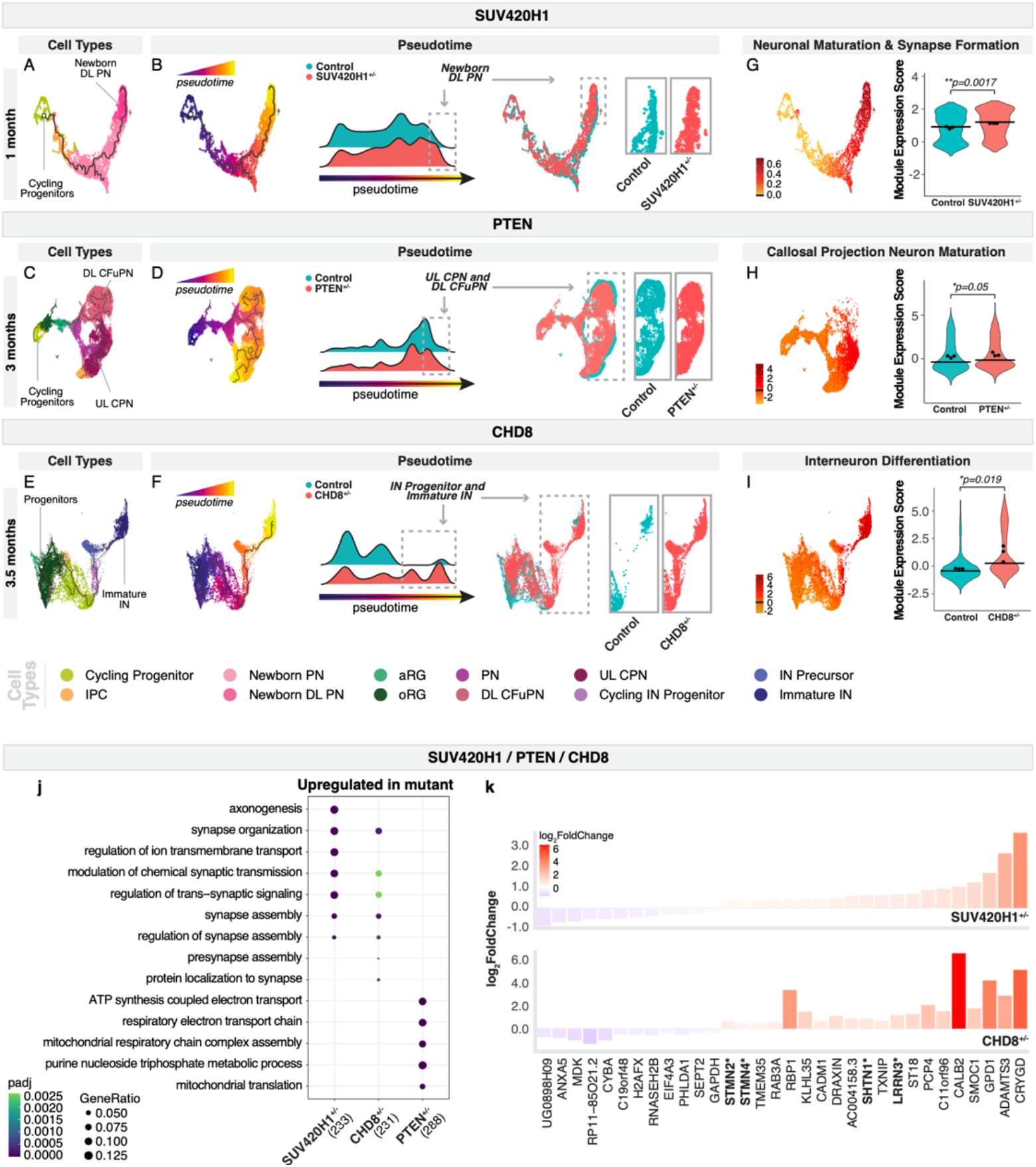
SUV420H1, PTEN, and CHD8 mutant organoids show accelerated differentiation of specific neuronal types. *A-F*, Pseudotime UMAPs for the 35 *d.i.v. SUV420H1*^+/−^ (*A, B*) and 3 month *PTEN*^+/−^ (*C, D*) and 3.5 month *CHD8*^+/−^ (*E, F*) organoids, and their corresponding controls. Cells are colored according to cell type (*A, C, E*), pseudotime (blue, early to yellow, late) (*B, D, F* Left), and genotype (control, blue; mutant, red) (*B, D, F* Right). n = 3 single organoids per batch per condition. For *SUV420H1* and *CHD8*, one partition is shown. For full trajectories, see Supplementary Figure 6*A* and Supplementary Figure 7*A*, *B*. All cells are shown for *PTEN*. *G, H, I*, Modules of highly correlated genes identified through co-expression network analysis on cells within the partitions of interest. Examples of modules that are differentially expressed between control and mutant organoids are shown; names were assigned to each module based on the known roles of the genes included. Left, module scores per cell are shown on associated UMAPs. Right, violin plots show the distribution of module scores between controls and mutants. Horizontal bars show median scores, and dots show average score per organoid. Adjusted *p*-values show differences between control and mutant based on linear mixed models (see Methods). Additional modules are shown in Supplementary Figure 6 and Supplementary Figure 7. *J*, Enriched gene ontology terms for genes upregulated in mutant vs. control. Genes were calculated using cells from the partitions of interest (i.e., the cells shown in g, h, and i). Size of dot indicates the proportion of genes belonging to each term found in the list of upregulated genes. Color indicates enrichment adjusted *p*-value. *K*, Gene expression changes of all genes significantly changing (adjusted *p* < 0.05) in the same direction in *SUV420H1*^+/−^ 35 *d.i.v.* and in *CHD8*^+/−^ 3.5 month cells shown in g, i. Genes marked in bold with an asterisk were also trending towards upregulation in *PTEN*^+/−^ three month callosal projection neuron lineage cells (adjusted *p* < 0.2). Color and bar size indicates log_2_ fold change. PN, projection neurons; aRG, apical radial glia; UL CPN, upper layer callosal projection neurons; oRG, outer radial glia; DL CFuPN, deep layer corticofugal projection neurons; DL, deep layers; IPC, intermediate progenitor cells; IN: interneurons; padj: adjusted *p*-value.

Strikingly, for each of the three genes, mutant organoids showed an increased distribution of cells toward the end point of these developmental trajectories, i.e., at a more advanced stage of development, compared to controls (*p* < 2.2e-16 in all cases, one-sided Kolmogorov–Smirnov test, Figure 5B, D, F). Combined with the increased number of neurons in the mutant organoids, these results further support an accelerated neurogenesis phenotype in each of the mutants.

In order to explore molecular differences that may underlie these phenotypes, we examined gene expression regulation across the partitions of interest using a co-expression network analysis. Using Weighted Correlation Network Analysis (WGCNA) (Langfelder & Horvath, 2008), we found modules of genes with highly correlated expression across cells in each dataset. Full lists of modules and their associated genes can be found in Supplementary Figure 6 and Supplementary Figure 7. For each mutation, at least one module was expressed most highly towards the end of the trajectory and was found to be significantly increased in expression in mutant organoids. In the *SUV420H1* 35 *d.i.v.* dataset, several modules were identified (Supplementary Figure 6B). One of them was significantly upregulated in mutant organoids compared to control (adjusted *p* = 0.0017, linear mixed models, Figure 5G) and contained multiple genes known to be involved in neuronal maturation and synapse formation (e.g., *GRIA2*, *CAMK2B*, *SNAP25*, *CNTNAP2*, *NRCAM*, and *NRNX2*).

For the three month *PTEN* mutant organoids, modules were separately calculated for each branch of the pseudotime trajectory (Supplementary Figure 7A): the upper layer callosal projection neuron and the deep layer corticofugal projection neuron branches (Supplementary Figure 7C, D). In the trajectory to upper layer callosal projection neurons (the partition of interest), several modules were found to be differentially expressed (Supplementary Figure 7C); in particular, a module containing genes related to callosal projection neurons and neuronal maturation (e.g., *SATB2*, *CAMKV*, *NELL2*, *SNX7*, and *SYBU*) showed increased expression in mutant cells (adjusted *p* = 0.05, linear mixed models, Figure 5H). Finally, in 3.5 month *CHD8* mutant organoids, a module containing genes related to interneuron differentiation (e.g., *DLX1*, *DLX2*, *DLX6-AS1*, *DLX5*, *SCGN*, and *GAD2*) was upregulated in the mutant (adjusted *p* = 0.019, linear mixed models, Figure 5I), while modules related to progenitor biology were downregulated (Supplementary Figure 7E).

These module analyses demonstrate altered gene regulation consistent with accelerated neuronal maturation. Collectively, these results indicate that mutations in *SUV420H1*, *PTEN*, and *CHD8* each lead to an acceleration of the developmental trajectory of specific classes of human cortical neurons.

### *CHD8* and *SUV420H1* haploinsufficiency share signatures of synaptic processes

It is conceivable that although affecting different neuron types, different ASD risk genes may act through common molecular pathways or mechanisms. To investigate this possibility, we compared differential gene regulation changes across the three ASD mutations. Reads from the cells in each selected partition of interest (Supplementary Fig 6A and Supplementary Figure 7A, B) were summed within each organoid, and DESeq2 (Love et al., 2014) was used to calculate differentially expressed genes (DEGs) between mutant and control in each dataset (a technique which explicitly controls for sample-to-sample variation (Lun & Marioni, 2017)). We did not observe overlap of GO terms for genes downregulated in any of the three mutations (Supplementary Figure 8A). Remarkably, synaptic gene ontology terms such as “synapse organization” and “regulation of trans-synaptic signaling” were significantly enriched (*p* < 0.0025 in all cases, over-representation test; Figure 5J) in genes upregulated in both *SUV420H1* (Figure 5A) and *CHD8* (Figure 5E) mutant organoids, despite affecting different neuronal cell types and time points. Although these GO terms were not significantly enriched for genes dysregulated in the *PTEN* dataset, some of the same genes were upregulated, or trending towards upregulation, in the trajectory branch containing callosal projection neurons in *PTEN* organoids as well (marked with a * in Figure 5K). The consistent upregulation of these gene sets indicates convergent dysregulation in molecular pathways relating to synapse formation and function in *CHD8* and *SUV420H1* mutants compared to controls (Figure 5J, K).

We next tested the hypothesis of a broader convergence of altered protein networks across mutations. We thus built an integrated protein interaction network using the DEPs across mutations, and found that DEPs from all three mutations populated multiple sub-clusters of the network, enriched for terms including “synapse”, “axon extension”, and “calcium signaling” (adjusted *p* ≤ 0.01 in all cases; Supplementary Figure 8B). However, when we calculated protein-protein interactome weighted distance between the protein sets, we found that the protein sets were not closer than would be expected by chance (p > 0.25 in all cases; Supplementary Figure 8C). This indicates that at this stage of development, although *SUV420H1*, *CHD8* and *PTEN* mutations all affect the expression of proteins involved in neuronal processes, they diverge in the use of specific protein effectors in the cell types they affect.

Collectively, these data show that *SUV420H1*, *CHD8*, and *PTEN* heterozygous organoids converge on a phenotype of shared accelerated cortical neurogenesis that is associated with the overexpression of genes implicated in synapse biology, but act on distinct neuronal types and without global convergence on the same molecular pathways.

## DISCUSSION

The process by which mutations in ASD risk genes converge on the neurobiology of this multifaceted disorder remains unclear. Progress in this field has been difficult because of the complexity of ASD genetics, limited experimental access to the human brain, and the lack of experimental models for human brain development (Velasco et al., 2020). By leveraging reproducible human cortical organoids, proteomics, and single cell genomics, we find that mutations in three ASD risk genes, *SUV420H1*, *PTEN*, and *CHD8* converge on a shared phenotype of accelerated early cortical neurogenesis, and define three neuronal classes of the cortical microcircuit as the specific cell types affected. These findings are interesting in light of prior work indicating that ASD genetics affects human neurogenesis (Adhya et al., 2020, Cederquist et al., 2020, Durak et al., 2016, Lalli et al., 2020, Lennox et al., 2020, Marchetto et al., 2017, Mariani et al., 2015, Wang et al., 2017, Zhang et al., 2020), and it underscores the value of organoids to identify both the developmental processes affected and the target cell types involved.

Our finding that different ASD risk genes converge in accelerated neuronal production but diverge one level of complexity below it, with each mutation affecting distinct neuronal types, suggests that the shared clinical pathology of these genes may derive from high-order processes of neuronal differentiation and circuit wiring, as opposed to convergent use of the same cellular and molecular mechanisms. Indeed, for the three ASD risk genes analyzed we found minimal sharing of molecular pathways. These results inform a framework for therapeutic approaches that might be better tailored to the modulation of shared dysfunctional circuit properties in addition to shared molecular pathways.

Excitatory/inhibitory imbalance of the cortical microcircuit is a major hypothesis for the etiology of ASD (Dani et al., 2005, Gogolla et al., 2009, Rubenstein & Merzenich, 2003). Such an imbalance could derive from a change in the numerical proportions of cell types within the circuit, incorrect wiring, or altered properties of the neurons and glial cells involved. The increased number of neurons that we observe in mutant organoids points to an early defect in circuit development whereby a small number of neuron classes develop asynchronously with the remaining cell types of the circuit. Studies *in vivo* suggest that these types of relatively subtle developmental alterations, even if resolved later in development, can result in disproportionally large functional effects on the mature circuit (Bocchi et al., 2017, Gao et al., 2013, Li et al., 2018, Lodato et al., 2011a, Lodato et al., 2011b, Smith & Fitzpatrick, 2012). Future work to increase the fidelity of circuit and tissue architecture in organoids will be crucial to investigate the implications of ASD risk alterations and potential interventions on circuit physiology and function.

Reproducible brain organoid models, combined with high-throughput single cell genomic methods, are suitable systems for unbiased identification of pathogenic mechanisms in neurodevelopmental disorders. This work paves the way for larger efforts to multiplex perturbation and analysis of large spectra of ASD risk genes (Jin et al., 2019) in organoids to understand whether convergent mechanisms are at play across the vast number of implicated genes, and for studies examining whether more complex polygenic disease states may also converge on similar shared mechanisms.

## CODE AND DATA AVAILABILITY

Scripts used for data analysis in this paper can be accessed at https://github.com/AmandaKedaigle/asd-mutated-brain-organoids.

Count data from all scRNA-seq experiments can be interactively explored at https://singlecell.broadinstitute.org/single_cell/study/SCP1129.

## SUPPLEMENTARY FIGURES

**Supplementary Figure 1.**
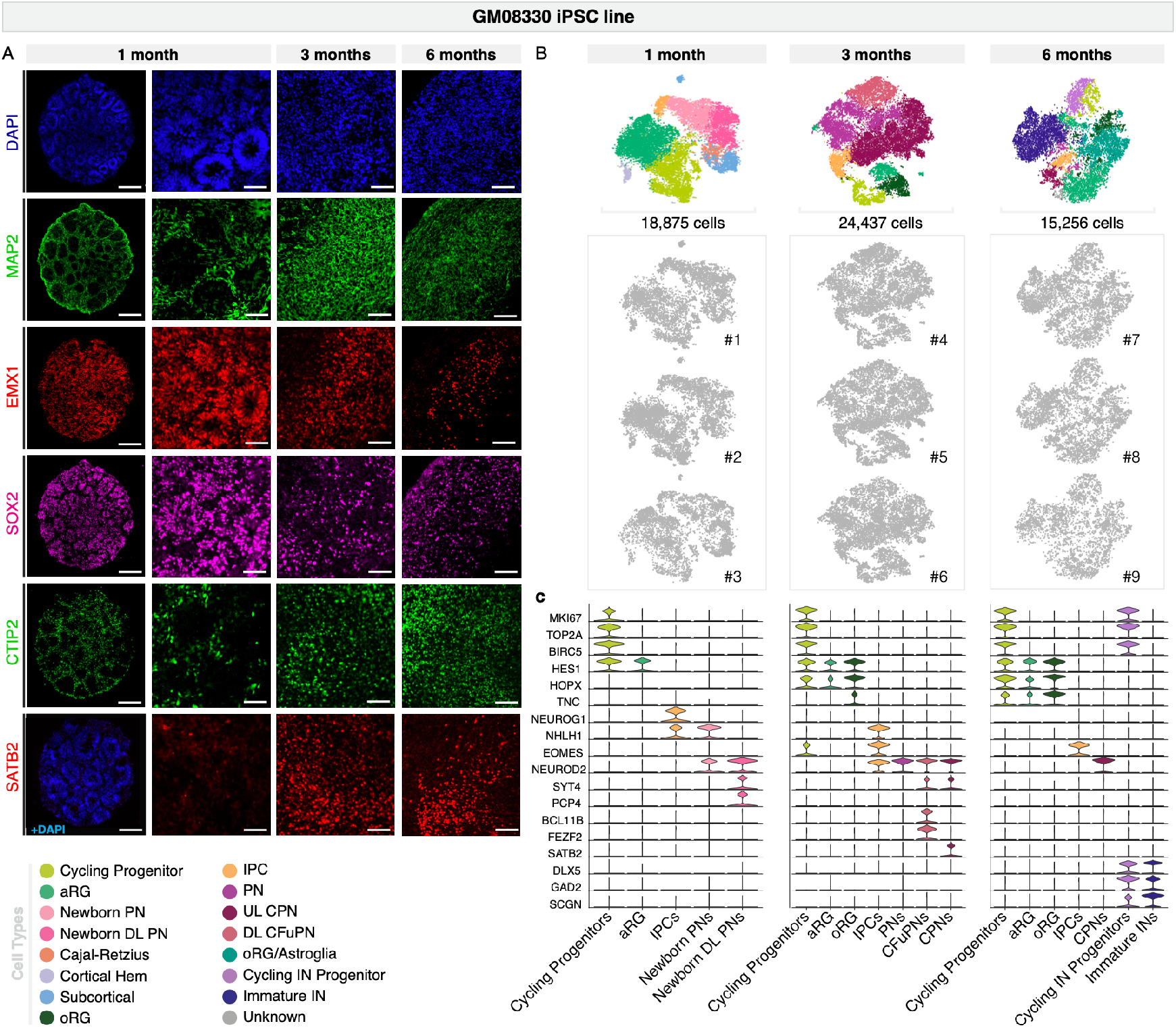
Cortical organoids cultured for one, three and six months generate the cellular diversity of the human cerebral cortex with high organoid-to-organoid reproducibility. *A*, Immunohistochemistry for neuronal (MAP2), dorsal forebrain neural progenitor (EMX1, SOX2), CFuPN (CTIP2), and CPN (SATB2) markers in GM08330 organoids at one, three, and six months. Scale bars: whole organoids, 200 μm; others, 50 μm. *B*, t-SNE plots of scRNA-seq data from individual one, three, and six months organoids from the GM08330 cell line. Top, t-SNE plots for the three organoids from each time point after Harmony batch correction, colored by cell types, repeated from Figure 1*A*, *B* and *C* for ease of comparison. *C*, Expression of selected marker genes used in cell-type identification. Violin plots show distribution of normalized expression in cells from organoids at one, three and six months (n = 3 individual organoids per time point). PN, projection neurons; aRG, apical radial glia; UL CPN, upper layer callosal projection neurons; oRG, outer radial glia; DL CFuPN, deep layer corticofugal projection neurons; DL, deep layers; IPC, intermediate progenitor cells; IN: interneurons; t-SNE, t-distributed stochastic neighbor embedding.

**Supplementary Figure 2.**
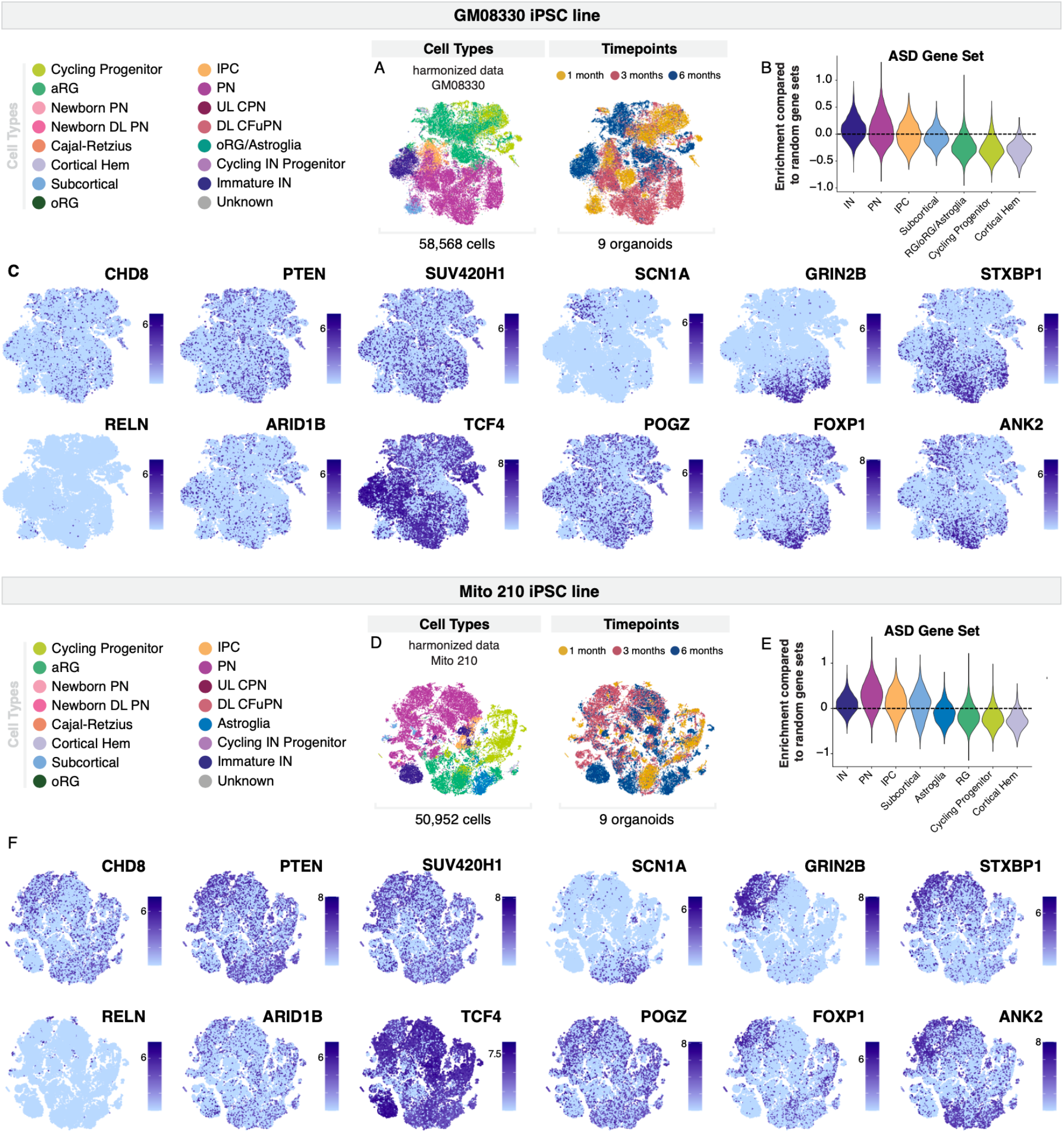
Expression of selected ASD risk genes in cortical organoids cultured for one, three, and six months from two cell lines. *A*, t-SNE plots of 58, 568 cells from 9 organoids from the GM08330 cell line, repeated from Figure 1*D*. Cells are colored according to cell type (left) or time point (right). *B*, Gene set scores for a set of 102 genes associated with ASD risk(Satterstrom et al., 2020) across cell types, in cells from a. Repeated from Figure 1*E* for ease of comparison. Scores above 0 indicate higher expression than similar modules of randomly chosen genes. *C*, t-SNE plots showing normalized expression of selected ASD risk genes in cells from a. *D*, t-SNE plots of 9 organoids from the Mito 210 cell line, at one, three and six months, after Harmony batch correction. Cells are colored according to cell type (left) or timepoint (right). *E*, Gene set scores for the set of ASD risk genes as in b, in cells from d. Scores above 0 indicate higher expression than similar modules of randomly chosen genes. *F*, t-SNE plots showing normalized expression of selected ASD risk genes in cells from d. PN, projection neurons; aRG, apical radial glia; UL CPN, upper layer callosal projection neurons; oRG, outer radial glia; DL CFuPN, deep layer corticofugal projection neurons; DL, deep layers; IPC, intermediate progenitor cells; IN: interneurons; t-SNE, t-distributed stochastic neighbor embedding.

**Supplementary Figure 3.**
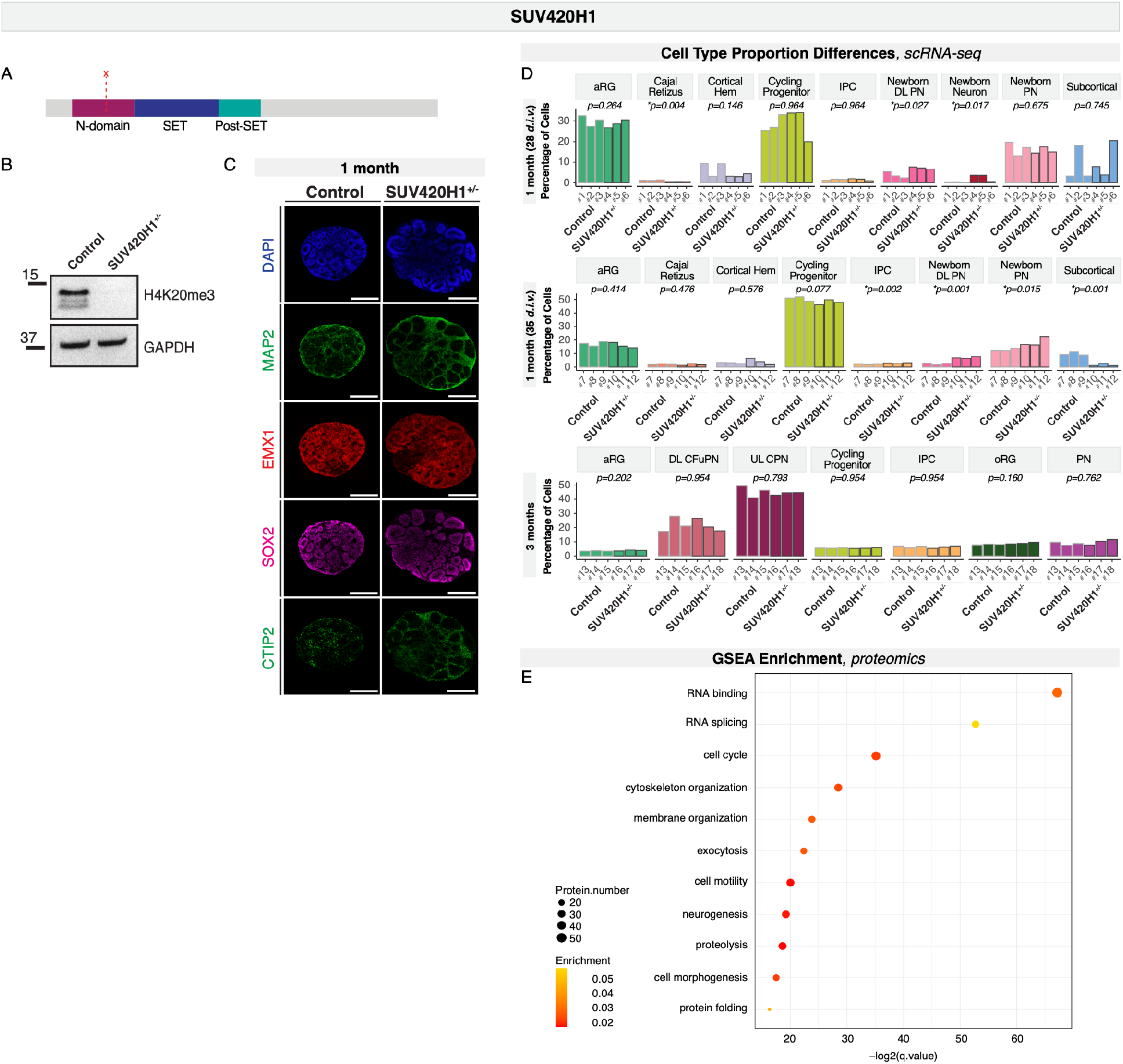
Validation of SUV420H1^+/−^ edited and isogenic control cell lines and generation of brain organoids. *A*, Protein domain structure of SUV420H1. In this study, the Mito 210 iPSC line was edited to induce a mutation in the N-domain (dashed line) that resulted in a protein-truncating variant. *B*, Western blot analysis showing presence of H4K20me3, a target of SUV420H1 activity, in control and high reduction in *SUV420H1*^+/−^ mutant organoids, using an antibody directed against this modification. *C*, Immunohistochemistry for neuronal (MAP2), dorsal forebrain neural progenitor (EMX1, SOX2) and CFuPN (CTIP2) markers in one month organoids derived from the Mito 210 *SUV420H1*^+/−^ mutant and isogenic control cell lines. Scale bar, 300 μm. *D*, Fraction of cells belonging to each cell type in individual organoids at 28 *d.i.v.*, 35 *d.i.v.*, and three months. Adjusted *p*-values show significance of change between mutant and control organoids, based on logistic mixed models (see Methods). *E*, Enriched gene ontology terms for DEPs between *SUV420H1*^+/−^ and control organoids cultured for 35 *d.i.v.* N-domain: N-terminal domain; H4K20me3, histone 4 lysine 20 trimethylation; *d.i.v*, days *in vitro*; PN, projection neurons; aRG, apical radial glia; UL CPN, upper layer callosal projection neurons; oRG, outer radial glia; DL CFuPN, deep layer corticofugal projection neurons; DL, deep layers; IPC, intermediate progenitor cells; DEPs: differentially expressed proteins.

**Supplementary Figure 4.**
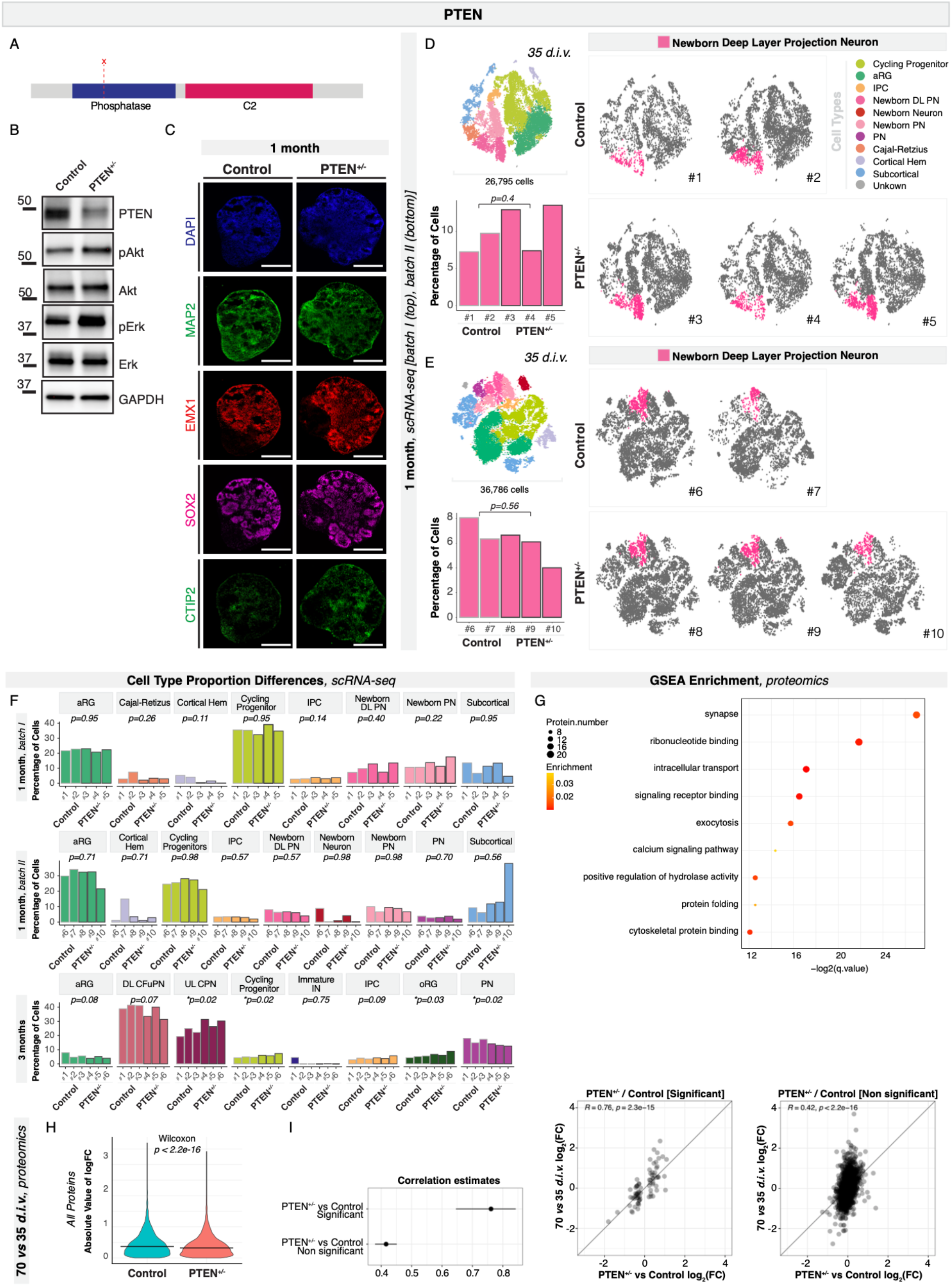
Validation of PTEN^+/−^ edited and isogenic control cell lines and generation of cortical organoids. *A*, Protein domain structure of PTEN. In this study, the Mito 210 iPSC line was edited to induce a mutation in the phosphatase domain (dashed line) that resulted in a protein-truncating variant. *B*, Western blot analysis showing presence of PTEN and its targets in control, and reduction of PTEN along with increase in phosphorylation of PTEN targets in mutant organoids, using an antibody directed against these modifications. *C*, Immunohistochemistry for neuronal (MAP2), dorsal forebrain neural progenitor (EMX1, SOX2), and CFuPN (CTIP2) markers in one month organoids derived from the Mito 210 *PTEN* mutant and isogenic control cell lines. Scale bar, 100 μm. *D*, *E*, scRNA-seq data from *PTEN*^+/−^ and control organoids cultured for 35 *d.i.v.*, batch I (*D*) and II (*E*). Top, Combined t-SNE plots of control and mutant organoids (n = 2 control and 3 mutant individual organoids). Cells are colored by cell type and the number of cells per plot is indicated. Right, t-SNE plots split into control and mutant organoids. Newborn deep layer projection neuron populations are highlighted in color. Bottom, bar charts showing the percentage of highlighted populations in each control and mutant organoid. Adjusted p values for a difference in cell type proportions between control and mutant, based on logistic mixed models (see Methods) are shown. *F*, Fraction of cells belonging to each cell type in individual organoids after one month (top and middle) and three months (bottom) *in vitro*. Adjusted *p*-values show significance of change between mutant and control organoids, based on logistic mixed models (see Methods). *G*, Enriched gene ontology terms for DEPs between *PTEN*^+/−^ and control organoids cultured for 35 *d.i.v. H*, Protein expression changes 70 vs. 35 *d.i.v.* for control and mutant organoids, for all detected proteins. *P*-value from a paired signed Wilcoxon rank test. *I*, Correlation between protein expression differences between *PTEN*^+/−^ and control organoids at 35 *d.i.v.*, and between control organoids at 70 vs. 35 *d.i.v.* Correlation is shown for proteins that significantly change between mutant and control organoids at 35 *d.i.v.* (middle), those that did not (right), and a comparison of their respective Pearson correlation coefficients (left). *d.i.v.*: days *in vitro;* PN, projection neurons; aRG, apical radial glia; UL CPN, upper layer callosal projection neurons; oRG, outer radial glia; DL CFuPN, deep layer corticofugal projection neurons; DL, deep layers; IPC, intermediate progenitor cell; DEPs: differentially expressed proteins; t-SNE, t-distributed stochastic neighbor embedding.

**Supplementary Figure 5.**
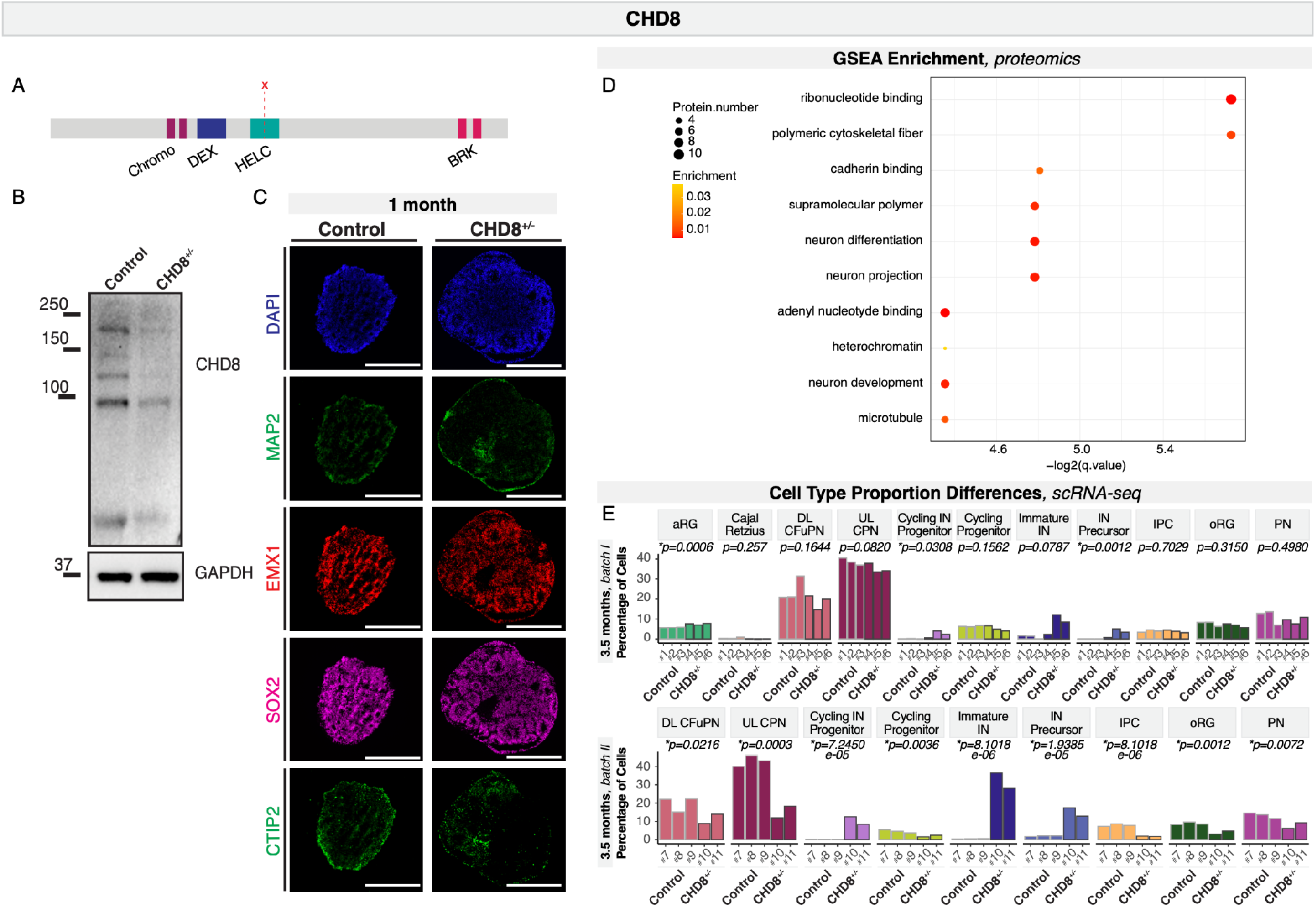
Validation of CHD8^+/−^ edited and isogenic control cell lines and generation of brain organoids. *A*, Protein domain structure of CHD8. The HUES66 line used in this study was engineered to carry a mutation in the helicase C-terminal domain (HELC domain, dashed line) that results in a protein-truncating variant. *B*, Western blot analysis showing presence of CHD8 in control, and reduction in mutant brain organoids, using an antibody directed against this protein. *C*, Immunohistochemistry for neuronal (MAP2), dorsal forebrain progenitor (EMX1, SOX2), and CFuPN (CTIP2) markers in one month organoids derived from the HUESS66 *CHD8* mutant and isogenic control cell lines. Scale bar, 100 μm. *D*, Enriched gene ontology terms for DEPs differentially expressed proteins between *CHD8*^+/−^ and control organoids cultured for 35 *d.i.v. E*, Fraction of cells belonging to each cell type in individual organoids. Adjusted *p*-values show significance of change between mutant and control organoids, based on logistic mixed models (see Methods). *d.i.v.*: days *in vitro;* PN, projection neurons; aRG, apical radial glia; UL CPN, upper layer callosal projection neurons; oRG, outer radial glia; DL CFuPN, deep layer corticofugal projection neurons; DL, deep layers; IPC, intermediate progenitor cell; IN, interneurons. DEPs: differentially expressed proteins.

**Supplementary Figure 6.**
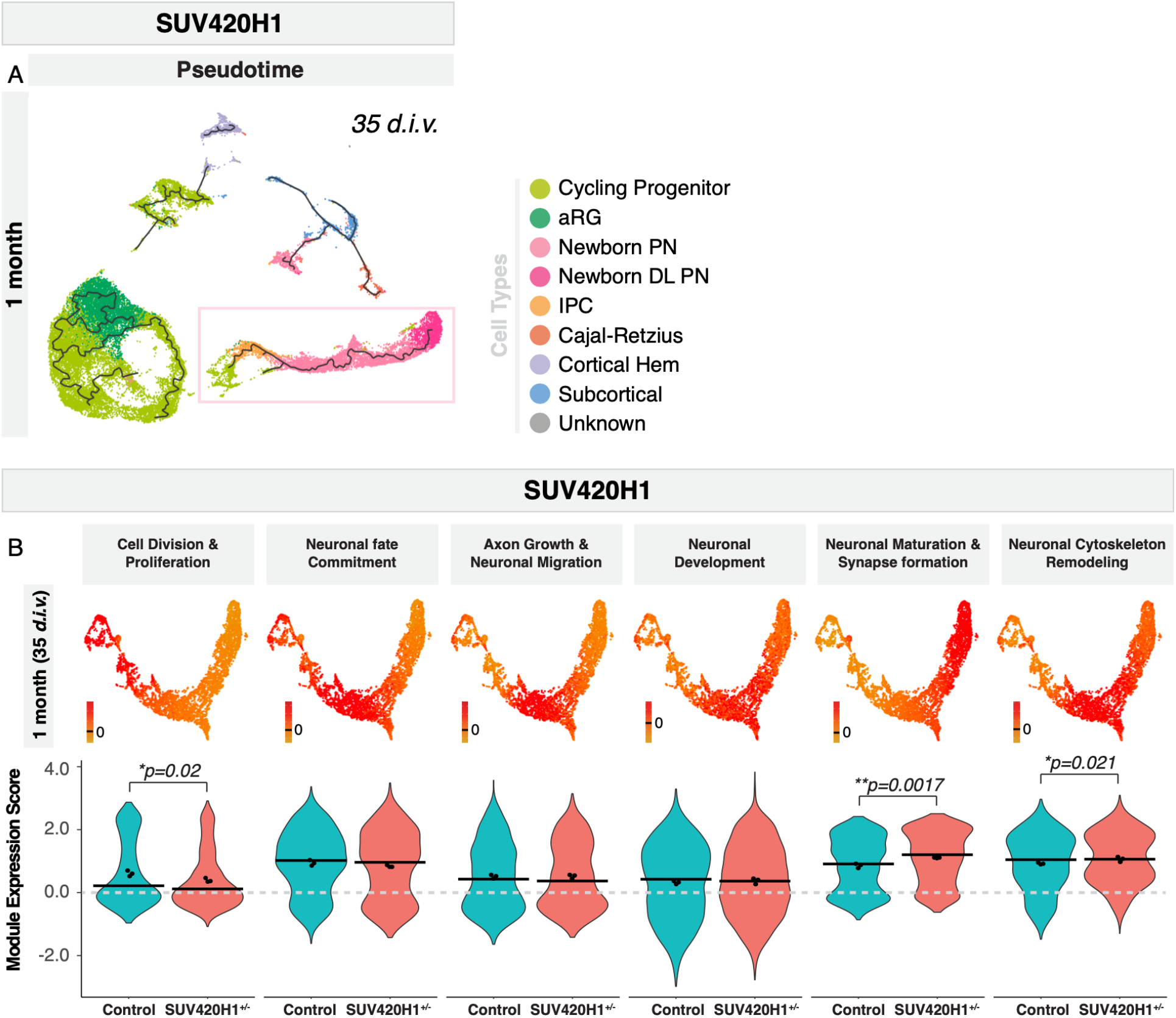
Full pseudotime trajectories and gene modules in SUV420H1^+/−^ and isogenic control organoids. *A-B*, Pseudotime trajectories from full datasets of *SUV420H1*^+/−^ 35 *d.i.v.* and control organoids, calculated with Monocle3. The partition highlighted by a box was subsetted and the trajectory is shown in Figure 5*A*. *B*, Module scores (top) and their distribution across mutant and control cells (bottom) for all modules resulting from WGCNA analysis of the partition of interest from *SUV420H1*^+/−^ and control organoids at 35 *d.i.v*. Cells were downsampled to have an equal number of cells per organoid. Names were assigned to each module based on the known functions of the genes included in each one. Horizontal bars show median scores, and dots show average score per organoid. Adjusted *p*-values show differences between control and mutant based on linear mixed models (see Methods). *d.i.v.*: days *in vitro;* PN, projection neurons; aRG, apical radial glia; UL CPN, upper layer callosal projection neurons; oRG, outer radial glia; DL CFuPN, deep layer corticofugal projection neurons; DL, deep layers; IPC, intermediate progenitor cell.

**Supplementary Figure 7.**
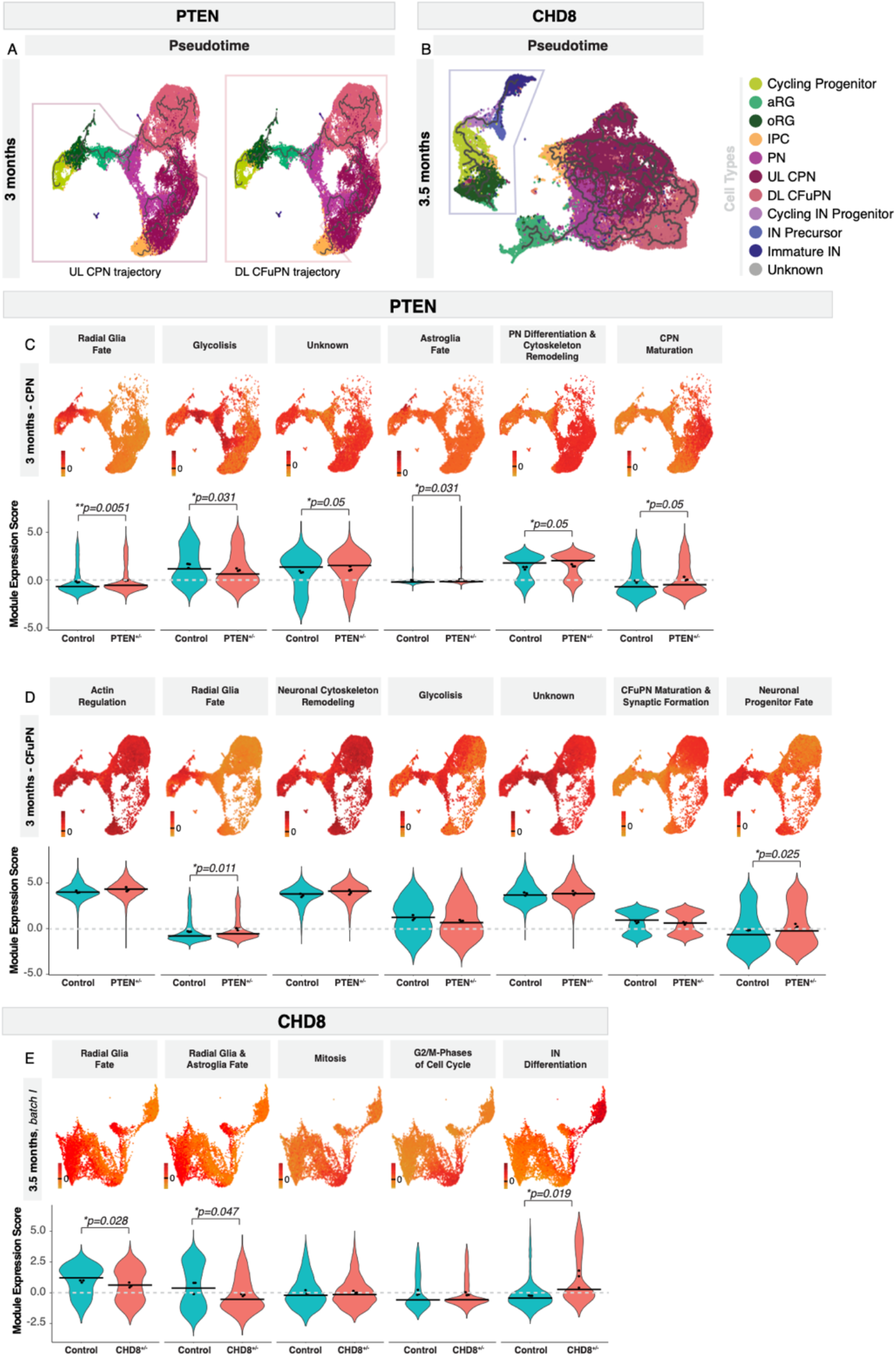
Full pseudotime trajectories and gene modules in PTEN^+/−^ and CHD8^+/−^ and isogenic control organoids. *A-B*, Pseudotime trajectories from full datasets of *PTEN*^+/−^ 3 months (*A*) and *CHD8*^+/−^ 3.5 months *in vitro* (*B*) and their control organoids, calculated with Monocle3. The partition highlighted by a box (left box of *A*) was subsetted and the trajectory is shown in Figure 5*E*. *C-E*, Module scores (top) and their distribution across mutant and control cells (bottom) for all modules resulting from WGCNA analysis of the partitions of interest. Cells were downsampled to have an equal number of cells per organoid. Names were assigned to each module based on the known functions of the genes included in each one. Horizontal bars show median scores, and dots show average score per organoid. Adjusted *p*-values show differences between control and mutant based on linear mixed models (see Methods). *d.i.v.*: days *in vitro;* PN, projection neurons; aRG, apical radial glia; UL CPN, upper layer callosal projection neurons; oRG, outer radial glia; DL CFuPN, deep layer corticofugal projection neurons; DL, deep layers; IPC, intermediate progenitor cell; IN, interneurons.

**Supplementary Figure 8.**
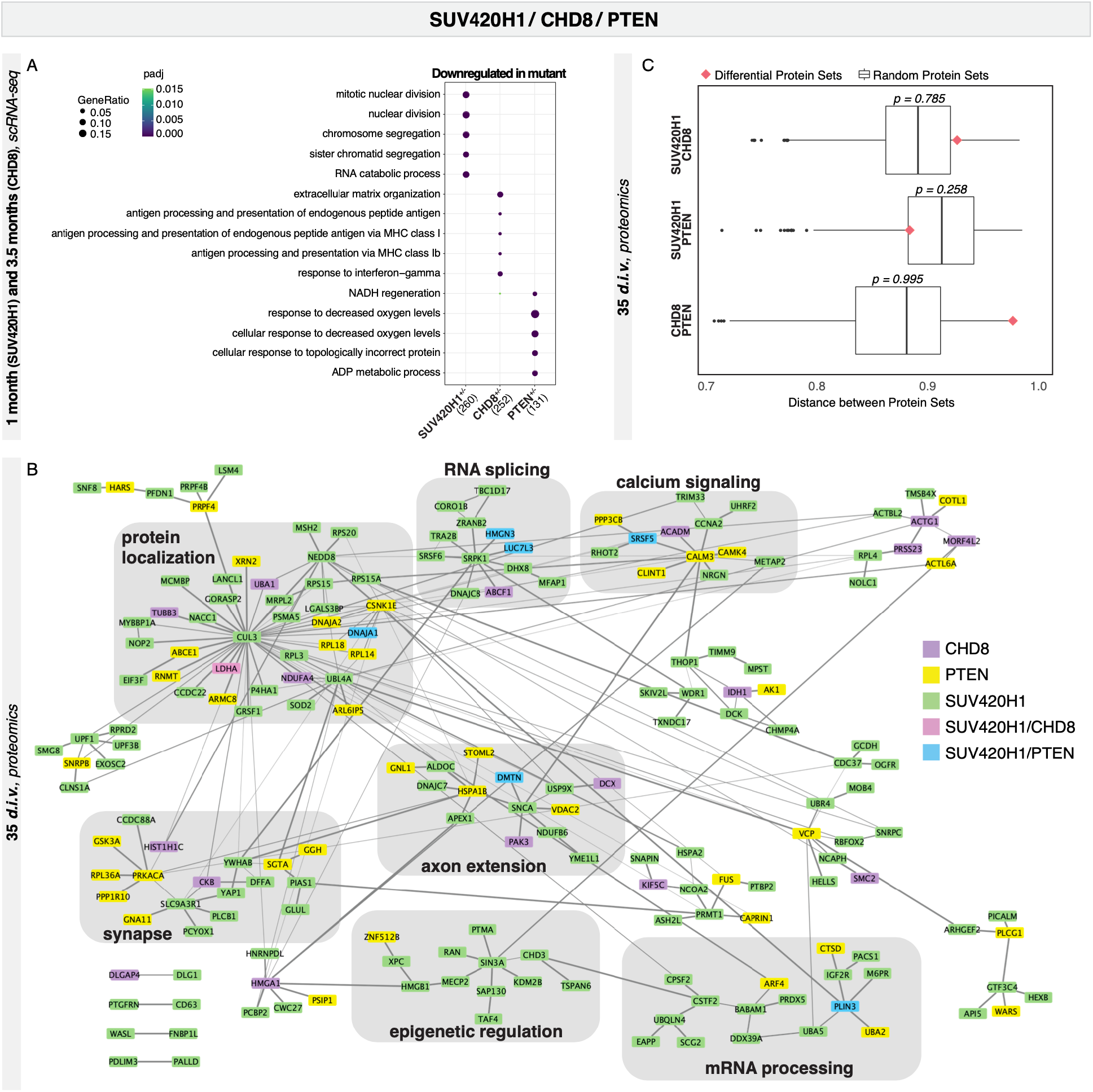
Biological processes affected in proteomics and scRNA-seq from the three mutations. *A*, Enriched gene ontology terms for genes downregulated in mutant vs. control cells in each of Figure 5*G*, *H*, *I*. Size of dot indicates the proportion of genes belonging to each term found in the list of upregulated genes. Color indicates adjusted *p*-value. *B*, Protein-protein interaction network using DEPs from all three sets of mutant organoids, created using the prize-collecting Steiner forest algorithm. Protein nodes are colored by the mutant(s) that they were differentially expressed in. Lines between nodes indicate physical protein-protein interactions from the STRING database, where line thickness correlates with interaction confidence. Selected subclusters of the network and significantly enriched terms for those subclusters are highlighted with gray rectangles and black text. *C*, Protein set distances between pairs of differentially expressed protein sets. For each pair of mutations, a PPI-weighted protein set distance (Yoon et al., 2019) was calculated between DEPs (pink diamond). In order to determine if this distance was smaller than would be expected by chance, size-matched sets were randomly chosen from the proteins detected in each experiment, and distance between these random sets was calculated 1000 times per pair. *P*-values were assigned by counting the number of times this random distance was less than the actual distance value between differential sets. DEPs: differentially expressed proteins.

## METHODS

### Pluripotent stem cell culture

The HUES66 ESC line(Chen et al., 2009) and mutant *CHD8* line (HUES66 AC2) were kindly provided by N. Sanjana, X. Shi, J. Pan, and F. Zhang (Broad Institute of MIT and Harvard); the Mito 210 human male psychiatric control iPSC line by B. Cohen Lab (McLean Hospital); and the GM08330 iPSC line (*a.k.a.* GM8330-8) by M. Talkowski Lab (MGH Hospital) and was originally from Coriell Institute. Cell lines were cultured as previously described(Velasco et al., 2019a, Velasco et al., 2019b). All human PSC cultures were maintained below passage 50, were negative for mycoplasma (assayed with MycoAlert PLUS Mycoplasma Detection Kit, Lonza), and karyotypically normal (G-banded karyotype test performed by WiCell Research Institute). The HUES66 line was authenticated using STR analysis completed by GlobalStem. The Mito 210 line was authenticated by genotyping analysis (Fluidigm FPV5 chip) performed by the Broad Institute Genomics Platform. The GM08330 line was not authenticated. All experiments involving human cells were performed according to ISSCR 2016 guidelines, and approved by the Harvard University IRB and ESCRO committees.

### CRISPR-mediated gene editing of *SUV420H1* and *PTEN*

The CRISPR guides for *SUV420H1* and *PTEN* were designed using the Benchling CRISPR Guide Design Tool (Benchling Biology Software). For *SUV420H1*, a guide was designed to target the N-terminal domain to create a protein truncation early in the translated protein in all known protein coding transcripts. The *PTEN* guide was chosen to induce a stop codon in the catalytic phosphatase domain.

### Organoid Differentiation

Dorsal forebrain-patterned organoids were generated as previously described(Velasco et al., 2019a, Velasco et al., 2019b).

### Immunohistochemistry

Samples were prepared as previously described(Velasco et al., 2019a) with slight modifications to cryosection thickness (14-18 *μ*m) and tissue fixation time.

### Whole-Organoid Imaging

Organoids were processed using the SHIELD protocol(Park et al., 2019). Briefly, organoids were fixed for 30 minutes in 4% paraformaldehyde (PFA) at room temperature (RT) and were then treated with 3% (wt/v) polyglycerol-3-polyglycidyl ether (P3PE) for 48 hours in ice cold 0.1 M phosphate buffer (pH 7.2) at 4° C then transferred to 0.3% P3PE in 0.1 M sodium carbonate (pH 10) for 24 hours at 37° C. Samples were rinsed and cleared in 0.2 M SDS in 50 mM phosphate buffered saline (PBS, pH 7.3) for 48 hours at 55° C. Organoids were stained with Syto16 (Thermo Fisher Scientific #S7578), anti-SOX2 and anti-TBR1 using the SmartLabel system (Lifecanvas). Tissues were washed extensively for 24 hours in PBS + 0.1% Triton-X-100 and antibodies were fixed to the tissue using a 4% PFA solution overnight at RT. Tissues were refractive index-matched in PROTOS solution (RI = 1.519) and imaged using a SmartSPIM axially-swept light-sheet microscope (LifeCanvas Technologies). 3D image datasets were acquired using a 15x 0.4 NA objective (ASI-Special Optics, #54-10-12). Optional sections from whole-organoid datasets are shown in Fig. 1a and Fig. 2a.

### Microscopy and Organoid Size Analysis

Images of organoids in culture were taken with an EVOS FL microscope (Invitrogen), Lionheart™ FX Automated Microscope (BioTek Instruments), or with a Zeiss Axio Imager.Z2 (Zeiss). Immunofluorescence images were acquired with the latter two and analyzed with the Gen5 (BioTek Instruments) or Zen Blue (Zeiss) image processing software. ImageJ(Schneider et al., 2012) was used to quantify organoid size. Area values were obtained by tracing individual organoids on ImageJ software which measured area pixels. Measurements were plotted as a ratio to the average value for control organoids of each experimental batch.

### Western blotting

Proteins were extracted from iPSC using N-PER™ Neuronal Protein Extraction Reagent (Thermo Fisher Scientific) supplemented with protease (cOmplete™ Mini Protease Inhibitor Cocktail, Roche) and phosphatase inhibitor (PhosSTOP, Sigma). Organoids were dissociated using a P1000 tip and lysates were centrifuged for 10 minutes at 13,500 rpm at 4° C. Supernatants were transferred to new tubes. Protein concentration was quantified using the Pierce™ BCA Protein Assay Kit (Thermo Fisher Scientific). 15-20 μg of protein lysates were separated on a NuPAGE™ 4-12%, Bis-Tris Gel (Invitrogen) or Mini-PROTEAN 4–15% Gels (Bio-Rad) and transferred onto PVDF membrane. Blots were blocked in 5% nonfat dry milk (Bio-Rad) and incubated with primary antibodies overnight. Afterward, the blots were washed and incubated at RT with secondary horseradish peroxidase-conjugated antibodies (Abcam) for 1 hour. Blots were developed using SuperSignal™ West Femto Maximum Sensitivity Substrate (Thermo Fisher Scientific) or ECL™ Prime Western Blotting System (Millipore), and ChemiDoc System (Bio-Rad).

### Cell lysis and filter-aided sample preparation (FASP) digestion for mass spectrometry

For *SUV420H1* and *PTEN* 35 *d.i.v.*, 4 mutant and 4 control organoids were used. For *CHD8* 35 *d.i.v.*, and for *PTEN* 70 *d.i.v.*, 3 mutant and 3 control organoids were used. Cells were placed into microTUBE-15 (Covaris) microtubes with TPP buffer (Covaris) and lysed using a Covaris S220 Focused-ultrasonicator instrument with 125 W power over 180 seconds at 10% max peak power. Upon precipitation with chloroform/methanol, extracted proteins were weighed and digested according to FASP protocol (100 μg for *PTEN* and *CHD8*; 150 μg for *SUV420H1*). Briefly, the 10K filter was washed with 100 μl of triethylammonium bicarbonate (TEAB). Each sample was added, centrifuged, and the supernatant discarded. Then, 100 μl of Tris(2-carboxyethyl) phosphine (TCEP) 37 was added for one hour, centrifuged, and the supernatant discarded. While shielding from light, 100 μl IAcNH2 was added for 1 hour followed by spinning and discarding the supernatant. Next, 150 μl 50 mM TEAB + Sequencing Grade Trypsin (Promega) was added and left overnight at 38° C, upon which the samples were centrifuged and supernatant collected. Lastly, 50 μl 50 mM TEAB was added to the samples, followed by spinning and supernatant collection. The samples were then transferred to HPLC.

### TMT Mass tagging protocol peptide labeling

The TMT (Tandem Mass Tag) Label Reagents were equilibrated to RT and resuspended in anhydrous acetonitrile or ethanol (for the 0.8 mg vials 41 μl were added, for the 5 mg vials 256 μl were added). The reagent was dissolved for 5 minutes with occasional vortexing. TMT Label Reagent (41 μl, equivalent to 0.8 mg) was added to each 100-150 μg sample. The reaction was incubated for one hour at RT. Reaction was quenched after adding 8 μl of 5% hydroxylamine to the sample and incubating for 15 minutes. Samples were combined, dried in a speedvac (Eppendorf) and stored at −80° C.

### Hi-pH separation and mass spectrometry analysis

Before submission to LC-MS/MS (Liquid Chromatography with tandem mass spectrometry), each experiment sample was separated on a Hi-pH column (Thermo Fisher Scientific) according to the vendor’s instructions. After separation into 40 fractions, each fraction was submitted for a single LC-MS/MS experiment performed on a Lumos Tribrid (Thermo Fisher Scientific) equipped with 3000 Ultima Dual nanoHPLC pump (Thermo Fisher Scientific). Peptides were separated onto a 150 μm inner diameter microcapillary trapping column packed first with approximately 3 cm of C18 Reprosil resin (5 μm, 100 Å, Dr. Maisch GmbH) followed by PharmaFluidics micropack analytical column 50 cm. Separation was achieved through applying a gradient from 5–27% ACN in 0.1% formic acid over 90 minutes at 200 nl per minute. Electrospray ionization was enabled through applying a voltage of 1.8 kV using a home-made electrode junction at the end of the microcapillary column and sprayed from stainless-steel tips (PepSep). The Lumos Orbitrap was operated in data-dependent mode for the MS methods. The MS survey scan was performed in the Orbitrap in the range of 400 –1,800 m/z at a resolution of 6 × 104, followed by the selection of the 20 most intense ions (TOP20) for CID-MS2 fragmentation in the Ion trap using a precursor isolation width window of 2 m/z, AGC setting of 10,000, and a maximum ion accumulation of 50 ms. Singly-charged ion species were not subjected to CID fragmentation. Normalized collision energy was set to 35 V and an activation time of 10 ms. Ions in a 10 ppm m/z window around ions selected for MS2 were excluded from further selection for fragmentation for 90 seconds. The same TOP20 ions were subjected to HCD MS2 event in Orbitrap part of the instrument. The fragment ion isolation width was set to 0.8 m/z, AGC was set to 50,000, the maximum ion time was 150 ms, normalized collision energy was set to 34 V and an activation time of 1 ms for each HCD MS2 scan.

### Mass spectrometry data generation

Raw data were submitted for analysis in Proteome Discoverer 2.4 (Thermo Fisher Scientific) software. Assignment of MS/MS spectra was performed using the Sequest HT algorithm by searching the data against a protein sequence database including all entries from the Human Uniprot database(Bairoch & Apweiler, 1999, Consortium, 2018) and other known contaminants such as human keratins and common lab contaminants. Sequest HT searches were performed using a 10 ppm precursor ion tolerance and requiring each peptides N-/C termini to adhere with Trypsin protease specificity, while allowing up to two missed cleavages. 16-plex TMTpro tags on peptide N termini and lysine residues (+304.207 Da) was set as static modifications while methionine oxidation (+15.99492 Da) was set as variable modification. A MS2 spectra assignment false discovery rate (FDR) of 1% on protein level was achieved by applying the target-decoy database search. Filtering was performed using a Percolator (64 bit version)(Käll et al., 2008). For quantification, a 0.02 m/z window centered on the theoretical m/z value of each of the 6 reporter ions and the intensity of the signal closest to the theoretical m/z value was recorded. Reporter ion intensities were exported in the result file of Proteome Discoverer 2.4 search engine as Excel tables. The total signal intensity across all peptides quantified was summed for each TMT channel, and all intensity values were normalized to account for potentially uneven TMT labeling and/or sample handling variance for each labeled channel.

### Mass spectrometry data analysis

Potential contaminants were filtered out and proteins supported by at least two unique peptides were used for further analysis. We kept proteins that were missing in at most one sample per condition. Data were transformed and normalized using variance stabilizing normalization using the DEP package of Bioconductor(Zhang et al., 2018). To perform statistical analysis, data were imputed for missing values using random draws from a Gaussian distribution with width 0.3 and a mean that is down-shifted from the sample mean by 1.8. To detect statistically significant differential protein abundance between conditions we performed a moderated t-test using the LIMMA package of Bioconductor(Ritchie et al., 2015), employing an FDR threshold of 0.1. GSEA was performed using the GSEA software(Subramanian et al., 2005). GO and KEGG pathway annotation were utilized to perform functional annotation of the significantly regulated proteins. GO terms and KEGG pathways with FDR q-values < 0.05 were considered statistically significant. For PTEN datasets, correlation between mutant effect (PTEN^+/−^ *vs* control at 35 *d.i.v.*) and time effect (70 *vs* 35 *d.i.v.* in control organoids) was calculated using Pearson correlation. For PTEN 70 *vs* 35 *d.i.v.*, changes in protein levels in mutant and control organoids were compared to one another with a signed paired Wilcoxin rank test, using stat_compare_means from the ggpubr R package (https://rpkgs.datanovia.com/ggpubr/).

To build protein interaction networks, we used the prize-collecting Steiner forest algorithm(Tuncbag et al., 2013, Tuncbag et al., 2016) using all three sets of DEPs as terminals, with the absolute value of their log fold change as prizes. This algorithm optimizes the network to include high-confidence protein interactions between protein nodes with large prizes. We used the PCSF R package v0.99.1(Akhmedov et al., 2017) to calculate networks, with the STRING database as a background protein-protein interactome(Szklarczyk et al., 2019), using parameters r = 0.1, w = 0.8, b = 15, and mu = 0.01. As is default in that package, the network was subclustered using the edge betweenness clustering algorithm from the igraph package, and functional enrichment is performed on each cluster using the ENRICHR API. The Cytoscape software version 3.6.1 was used for network visualization(Shannon et al., 2003). To assess relationships between the three sets of differential proteins, a PPI-weighted gene distance(Yoon et al., 2019), was calculated between each pair of protein sets. A background distribution was calculated by drawing size-matched random lists of proteins from all detected proteins in each dataset and calculating the pMM between these sets. This was repeated 1000 times, and a *p*-value was calculated by evaluating the amount of times randomized pMMs were lower than the value calculated using DEPs.

### Dissociation of brain organoids and scRNA-seq

Organoids were dissociated as previously described(Arlotta et al., 2017, Velasco et al., 2019b) with minor adaptations of volumes for one month organoids. Cells were loaded onto a Chromium™ Single Cell B Chip (10x Genomics, PN-1000153), and processed them through the Chromium Controller to generate single cell GEMs (Gel Beads in Emulsion). scRNA-seq libraries were generated with the Chromium™ Single Cell 3’ Library & Gel Bead Kit v3 (10x Genomics, PN-1000075), with the exception of a few libraries in the earlier experiments that were prepared with a v2 kit (10x Genomics, PN-120237). We pooled libraries from different samples based on molar concentrations and sequenced them on a NextSeq 500 or NovaSeq instrument (Illumina) with 28 bases for read 1 (26 bases for v2 libraries), 55 bases for read 2 (57 bases for v2 libraries) and 8 bases for Index 1. If necessary, after the first round of sequencing, we re-pooled libraries based on the actual number of cells in each and re-sequenced with the goal of producing approximately equal number of reads per cell for each sample.

### scRNA-seq data analysis

Reads from scRNA-seq were aligned to the GRCh38 human reference genome and the cell-by-gene count matrices were produced with the Cell Ranger pipeline (10x Genomics)(Zheng et al., 2017). Cellranger version 2.0.1 was used for experiments using the GM08330 line and for CHD8 mutant and control organoids at 3.5 months - batch I, while version 3.0.2 was used for all other experiments. Default parameters were used, except for the ‘–cells’ argument. Data was analyzed using the Seurat R package v3.1.5(Stuart et al., 2019) using R v3.6. Cells expressing a minimum of 500 genes were kept, and UMI counts were normalized for each cell by the total expression, multiplied by 10^6^, and log-transformed. Variable genes were found using the “mean.var.plot” method, and the ScaleData function was used to regress out variation due to differences in total UMIs per cell. Principal component analysis (PCA) was performed on the scaled data for the variable genes, and top principal components were chosen based on Seurat’s ElbowPlots (at least 15 PCs were used in all cases). Cells were clustered in PCA space using Seurat’s FindNeighbors on top principal components, followed by FindClusters with resolution = 1.0. Variation in the cells was visualized by t-SNE (t-distributed stochastic neighbor embedding) on the top principal components.

In the case of GM08330 one month organoids, cells were demultiplexed using genotype clustering from cells from a different experiment that were sequenced in the same lane. To demultiplex, SNPs were called from CellRanger BAM files with the cellSNP tool v0.1.5, and then the vireo function was used with default parameters and n_donor = 2, from the cardelino R library v0.4.0(Huang et al., 2019, McCarthy et al., 2020) to assign cells to each genotype.

In three cases, one out of six organoids were excluded from the analysis as outliers. Specifically, in *PTEN*^+/−^ 35 *d.i.v.* batch I, a control organoid was excluded because not enough cells could be recovered from that 10x channel, and in the second batch, a control organoid was excluded because it clustered entirely separately from the other 5 organoids, indicating a potential differentiation issue. Finally, in *CHD8*^+/−^ 3.5 month batch II, a mutant organoid was excluded due to mostly containing interneuron lineage cells, with very few projection neuron cells. Although an increase in interneuron-lineage cells was seen in all mutant organoids in this experiment, this extreme example was excluded to be conservative. This left a total of 57 single organoids that passed quality control and were considered in downstream analysis, with a total of 416,051 cells.

For each dataset, upregulated genes in each cluster were identified using the VeniceMarker tool from the Signac package v0.0.7 from BioTuring (https://github.com/bioturing/signac). Cell types were assigned to each cluster by looking at the top most significant upregulated genes. In a few cases, clusters were further subclustered to assign identities at higher resolution.

To assess gene expression of ASD risk genes in GM08330 and Mito 210 control organoids across timepoints, datasets from one, three, and six months were merged using Seurat, then batch corrected using Harmony v1.0 with default parameters(Korsunsky et al., 2019). Since the one month data is dominated by cell cycle signal, the ScaleData function was used to regress out variation due to both total UMI count per cell and to cell cycle stage differences, calculated using Seurat’s CellCycleScore. Variation was visualized using t-SNE on the first 30 Harmony dimensions. Expression of the 102 ASD risk genes identified in the Satterstrom et. al. study(Satterstrom et al., 2020), compared to control gene lists with similar average expression, was evaluated using Seurat’s AddModuleScore function.

To compare cell type proportions between control and mutant organoids, for each cell type present in a dataset, the glmer function from the R package lme4 v1.1-23(Bates et al., 2015) was used to estimate a mixed-effect logistic regression model. The output was a binary indicator of whether cells belong to this cell type, the control or mutant state of the cell was a fixed predictor, and the organoid that the cell belonged to was a random intercept. Another model was fit without the control-versus-mutant predictor, and the anova function was used to compare the two model fits. *P-*values for each cell type were then adjusted for multiple hypothesis testing using the Benjamini-Hochberg correction.

### Pseudotime, gene module, and differential expression analysis

Pseudotime analysis was performed using the Monocle3 v. 0.2.0 software package(Cao et al., 2019) with default parameters. The cells were first subsetted to contain an equal amount from control and mutant. A starting point for the trajectory was chosen manually by finding an endpoint of the tree located in the earliest developmental cell type (generally, cycling progenitors). Where the cells were split into more than one partition, the starting point was chosen within the partition of interest, and a new UMAP was calculated using just these cells. To test whether mutant trajectories were accelerated compared to control, a one-sided Kolmogorov–Smirnov test was applied comparing the distribution of psuedotime values of control vs. mutant cells, using the stats R package.

In order to learn patterns of coordinated gene regulation across the cells, we applied WGCNA(Langfelder & Horvath, 2008) to each dataset. Where cells were split into partitions in the above pseudotime analysis, only cells belonging to the partition of interest were used. For the *PTEN* three months dataset, cells were split into two partitions based on the branching of the pseudotime into two subtypes of projection neurons: one excluding callosal projection neurons, and the other excluding corticofugal neurons. Normalized gene expression data was further filtered to remove outlying genes, mitochondrial and ribosomal genes. Outliers were identified by setting the upper (> 9) and lower (< 0.15) thresholds to the average normalized expression per gene. After processing, blockwiseModules function from the WGCNA v1.69 library was performed in R with the parameters networkType=”signed”, minModuleSize =4, corType =“Bicor”, maxPOutliers=0.1, deepSplit=3,trapErrors=T, and randomSeed=59069. Other than power, remaining parameters were left as the default setting. To pick an adequate power for each dataset, we used the pickSoftThreshold function from WGCNA to test values from 1 to 30. Final resolution was determined by choosing the resolution that captured most variation in the fewest total number of modules - this resulted in a power of 3 for *SUV420H1* 35 *d.i.v.*, and 12 for both partitions of *PTEN* 3 months and for *CHD8* 3.5 months.

To calculate differential expression of modules, Seurat objects were downsampled to have an equal number of cells per organoid, and then the AddModuleScore function was used, using gene lists from WGCNA results. For each module, linear mixed-effect models were fit to the data, with the modules scores as the output, the organoid the cell belongs to as a random intercept, and with or without the control-versus-mutant state as a predictor. The anova function was used to compare the models, and *p*-values were then adjusted across modules using the Benjamini-Hochberg correction.

Differential genes between control and mutant organoids were assessed after datasets were subsetted to the cells from the partition of interest in the above pseudotime analysis. Reads were then summed across cells in each organoid. Genes with less than 10 total reads were excluded, and DESeq2(Love et al., 2014) was used to calculate DEGs, with each organoid as a sample(Lun & Marioni, 2017). The clusterProfiler(Yu et al., 2012) R package was used to find enriched biological processes in these gene sets, with the enrichGO function and the compareCluster function to highlight processes the gene sets might have in common. Redundant GO terms were removed with the simplify function, with cutoff = 0.7.

## ACKNOWLEDGEMENTS

We thank J. R. Brown (from the P.A. laboratory) for editing the manuscript and the entire Arlotta Lab for support and insightful discussions; M. Daly, E. Robinson and B. Neale for insightful discussions on autism genetics; T. Nguyen (from G.Q. laboratory) for outstanding support with data analysis; X. Jin (from the P.A. laboratory) for helping with designing and sequencing of edited lines; A. Shetty (from the P.A. laboratory) for help with scRNA-seq cell type classification; N. Sanjana, X. Shi, J. Pan, and F. Zhang for the HUES66 AC2 line; the Talkowski laboratory for the GM08330 line; the Cohen laboratory for the Mito 210 line; the Ricci laboratory for providing nanoblades for the generation of SUV420H1 edited cell lines; L. Barrett at the Broad Institute Stem Cell Facility for generating the PTEN mutant line; L.M. Daheron at the Harvard Stem Cell Facility for expanding edited lines; and B. Budnik at the Harvard Center for Mass Spectrometry for conducting proteomics experiments. This work was supported by grants from the Stanley Center for Psychiatric Research, the Broad Institute of MIT and Harvard, the National Institutes of Health (R01-MH112940 to J.Z.L. and P.A., P50MH094271, and U01MH115727 to P.A.), the Klarman Cell Observatory to J.Z.L. and A.R., and the Howard Hughes Medical Institute to A.R.

## AUTHOR CONTRIBUTIONS

P.A., J.Z.L., B.P., S.V., A.J.K., M.P., G.Q. conceived all the experiments. A.D. designed the SUV420H1 and PTEN edited lines and generated the SUV420H1 line with help from B.P.. S.V., B.P., M.P., R.S., C.A., A.T, S.S. generated, cultured and characterized all the organoids in this study and P.A. supervised their work and contributed to data interpretation. X.A. performed scRNA-seq experiments, with help from B.P., S.V., R.S., G.Q.; A.J.K., K.K., S.K.S. and J.Z.L. performed scRNA-seq analysis and J.Z.L. and A.R. supervised the computational work. S.V., B.P., M.P., A.U., G.Q. and A.J.K. worked on cell type assignments and data analysis and P.A. supervised the work. K.T., M.P., A.J.K. performed proteomics analysis, supervised by K.L.. A.A. performed and processed whole-organoid imaging under the supervision of K.C.. P.A., B.P., S.V., A.J.K., and M.P. wrote the manuscript with contributions from all authors. All authors read and approved the final manuscript.

